# Specification and epigenetic resetting of the pig germline exhibit conservation with the human lineage

**DOI:** 10.1101/2020.08.07.241075

**Authors:** Qifan Zhu, Fei Sang, Sarah Withey, Walfred Tang, Sabine Dietmann, Doris Klisch, Priscila Ramos-Ibeas, Haixin Zhang, Cristina E. Requena, Petra Hajkova, Matt Loose, M. Azim Surani, Ramiro Alberio

## Abstract

Investigations on the human germline and programming are challenging due to limited access to embryonic material. However, the pig as a model may provide insight on transcriptional network and epigenetic reprogramming applicable to both species. Here we show that during the pre- and early migratory stages pig primordial germ cells (PGCs) initiate large-scale epigenetic reprogramming, including DNA demethylation involving TET-mediated hydroxylation and potentially base excision repair (BER). There is also macroH2A1 depletion and increased H3K27me3, as well as X chromosome reactivation (XCR) in females. Concomitantly, there is dampening of glycolytic metabolism genes and re-expression of some pluripotency genes like those in preimplantation embryos. We identified evolutionarily young transposable elements and gene coding regions resistant to DNA demethylation in acutely hypomethylated gonadal PGCs, with potential for transgenerational epigenetic inheritance. Detailed insights into the pig germline will likely contribute significantly to advances in human germline biology, including *in vitro* gametogenesis.

## Introduction

The germline transmits hereditary information, which ensures the continuity of the species. Development of the primordial germ cells (PGCs), the precursors of gametes, begins in peri-gastrulation embryos and is governed by a network of transcriptional regulators. Extensive epigenetic reprogramming follows, which includes the erasure of imprints and potentially epimutations for the restoration of totipotency (Hill et al., 2018; Kurimoto et al., 2008; Tang et al., 2016). While the principles of mammalian germline development are emerging, so are some important differences and gaps in our knowledge (Kobayashi and Surani, 2018; Saitou and Miyauchi, 2016).

We previously showed that the molecular program of pig PGCs (pPGCs) corresponds to what is known about human PGCs (hPGCs), indicating that studies in the pig may be informative for understanding the development of hPGCs (Kobayashi et al., 2017). A critical period of human germline development is between week (Wk) 2 and Wk 4, when PGCs are specified and migrate towards the gonads (Leitch et al., 2013). However, human embryos are not accessible during these critical stages, consequently, we have little or no information of germline development during this period.

At the equivalent developmental period in pigs, pPGCs are specified between embryonic (E) days 12-14, following sequential upregulation of SOX17 and BLIMP1 in response to BMP signalling (Kobayashi et al., 2017), as is the case during the induction of human PGC-like cells (hPGCLCs) *in vitro* (Irie et al., 2015). In pigs, pPGCs commence migration at ∼E15 through the hindgut, until reaching the gonadal ridges by E22, and undergo extensive proliferation between E28-42 (Hyldig et al., 2011a; Hyldig et al., 2011b).

Shortly after pPGC specification, pre-migratory pPGCs display initiation of epigenetic reprogramming, characterized by global reduction in DNA methylation and H3K9me2 (Hyldig et al., 2011a; Kobayashi et al., 2017; Petkov et al., 2009). Upon colonization of the gonads, pPGCs show asynchronous demethylation of imprinted genes and retrotransposons (Hyldig et al., 2011a; Hyldig et al., 2011b; Petkov et al., 2009). Accordingly, there is protracted epigenetic reprogramming in the pig germline over a period of several weeks.

Studies on early human PGCs have relied on pluripotent stem cell-based *in vitro* models, which showed that hPGCLCs originate from cells with posterior PS/incipient mesoderm-like identity following exposure to BMP, revealing SOX17 to be a critical determinant of the PGC fate (Irie et al., 2015; Kojima et al., 2017). Studies on *ex vivo* hPGCs showed that epigenetic reprogramming in the human germline is also protracted and asynchronous compared to mice (Gkountela et al., 2015; Guo et al., 2015; Tang et al., 2015), but there is limited scope for detailed investigations on *ex vivo* human embryos. We posit that investigations in the pig that develops as bilaminar discs, unlike egg cylinders of laboratory rodents, might provide insights into fundamental mechanisms of germline development that would apply widely to non-rodents, including the human germline.

Here, using single cell transcriptome (scRNASeq) and whole-genome bisulfite sequencing (WGBS), we reveal the transcriptional program and epigenetic features of pPGCs during a critical interval of development that is largely inaccessible for humans. We observed a close transcriptional alignment between pig and human PGCs. We also observed extensive epigenetic reprogramming characterized by DNA demethylation, XCR and histone modifications in pre- and early migratory pPGCs. A metabolic dampening of glycolytic metabolism genes and the reactivation of some pluripotency-associated genes accompanied these events. We identified genomic loci escaping global DNA demethylation, with potential for transgenerational epigenetic inheritance.

## Results and Discussion

### A highly conserved transcriptional program of pPGCs and hPGCs

Pig PGCs first emerge in E12 embryos, forming a cluster of ∼60 cells that expands to ∼150-200 by E14 (Kobayashi et al., 2017). To investigate the transcriptome of pre-migratory pPGCs, we dissected the posterior region of E14 embryos. We also isolated germ cells from E31 gonads (Table S1). After dissociation of the tissues into single cells and FACS sorting using an anti-Sda/GM2 antibody (Klisch et al., 2011), we manually picked individual cells for analysis (Fig. 1A and Fig. S1A). We obtained single-cell RNASeq (scRNASeq) data of 17 Sda/GM2^+^ cells (pre-migratory pPGCs) and 89 Sda/GM2^-^ (surrounding cells) from E14 embryos. We similarly analysed 22 Sda/GM2^+^ early (E31) gonadal PGCs using the Smart-Seq2 protocol (Picelli et al., 2014). After sequencing we identified closely related cells using unsupervised hierarchical clustering (UHC) and t-stochastic neighbour embedding (t-SNE) analysis, including a dataset of pig E11 epiblast (Ramos-Ibeas et al., 2019) (Fig. 1B and 1C). Epiblast (Epi) cells and E14 surrounding somatic cells cluster separately from pPGCs (Fig. 1B, and 1C). In both E14 and E31 pPGCs, we detected *PRDM1 (BLIMP1), TFAP2C, NANOS3* and *KIT*, and high expression of pluripotency genes *NANOG* and *POU5F1*. The late PGC markers *DAZL, DDX4* and *PIWIL2* were only detected in E31 gonadal PGCs. We did not detect *SOX2* in most pPGCs, which is also the case in hPGCs. We found expression of *PDPN, HERC5 and MKRN1* (Fig. 1B), which was recently reported in early hPGCs from a rare gastrulating Carnegie Stage 7 human embryo (CS7 hPGCs)(Tyser et al., 2020). SOX17 protein was present in pre-migratory and gonadal pPGCs observed by immunofluorescence (IF) (Fig. 1A and Fig. S1A), although the *SOX17* transcript was found in a subset of pPGCs (6/17 in E14 and 12/22 in E31)(Fig. 1B). Interestingly, low and fluctuating *SOX17* expression is also observed in early hPGCs in CS7 human embryo, while the endoderm lineage shows consistent and high SOX17. The low and fluctuating SOX17 expression in early pPGCs and hPGCs might reflect a conserved mechanism to regulate gene dosage to prevent expression of endoderm genes in human and pig PGCs (Irie et al., 2015; Kobayashi et al., 2017; Tyser et al., 2020).

**Figure 1.**
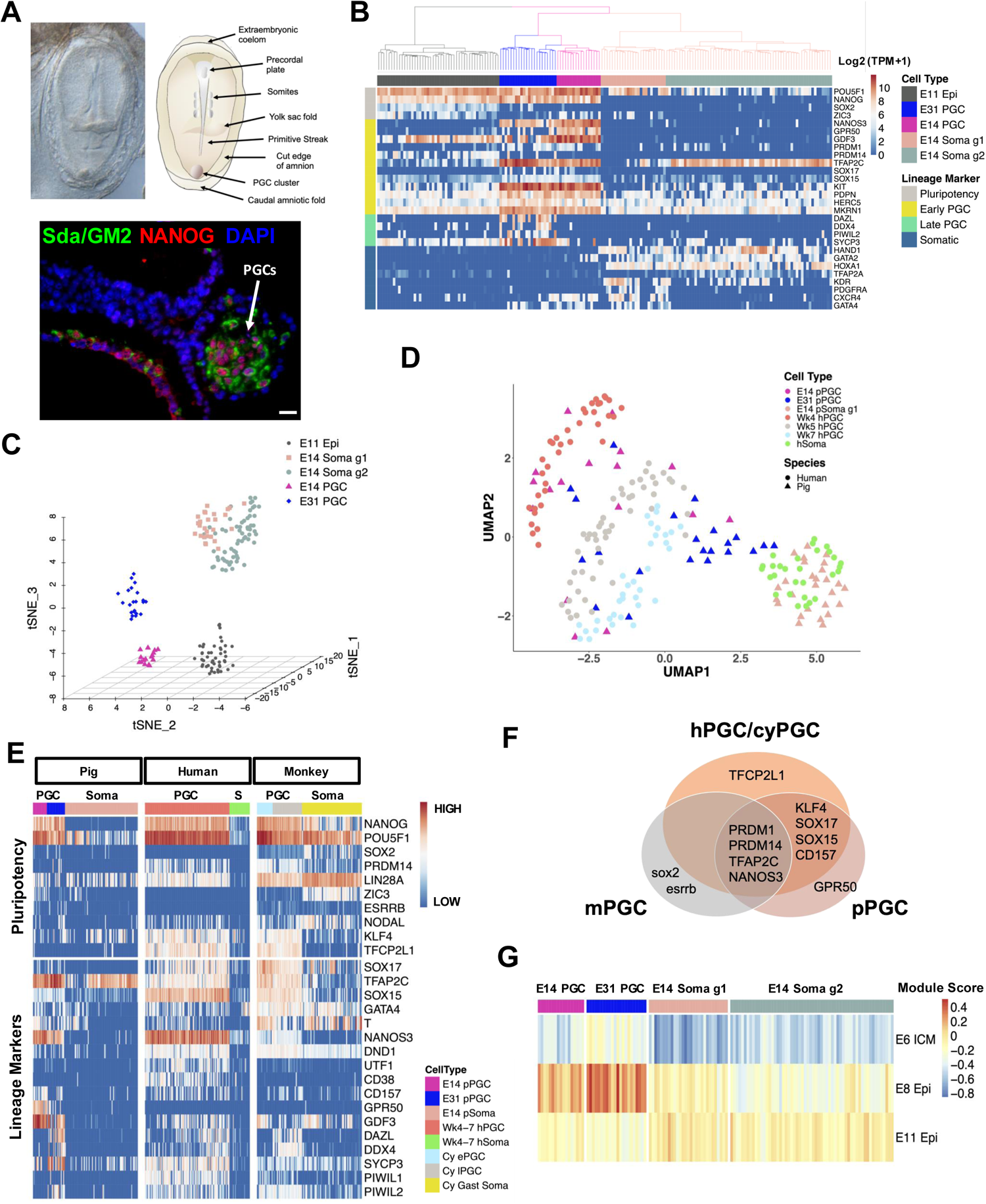
Transcriptional profile of pig PGC and comparison with hPGCs. (A) Isolation of pre-migratory (Pre-migr.) PGCs from E14 embryos with FACS sorting using the cell surface marker Sda/GM2. Immunofluorescence image of a midline sagittal section of an E14 embryo shows PGC cluster (white arrow) in the caudal end. Scale bar: 20µm. (B) Unsupervised hierarchical clustering of all expressed genes. A subset of known marker genes was selected for the heatmap.) and (C) t-SNE showing relationships between E11 Epiblast cells, E14 Somatic cells, E14 PGC and E31 PGC. (D) UMAP plot showing integration of human (Wk4-7) and pig (E14-31) PGCs and soma. (E) Expression profiles of pluripotency genes and lineage markers in pig, human and cyPGCs and somatic cells. “Cy ePGC”: early cyPGC (E13-20); “Cy lPGC”: late cyPGC (E36-55); “Cy Gast Soma”: cy gastrulating cells (E13-20). As different expression units are used for the three species, values in the colour scale are replaced by HIGH and LOW. (F) Schematic of unique or common genes between pig, human and mouse PGC. (G) Relative average expression (module score) of signature sets of E6, E8 and E11 cells in E14 and E31 cells. See also Figure S1, Figure S2, Table S1, and Table S4.

The posterior somatic cells in E14 embryos, which are the likely neighbours of pPGCs, segregated into two clusters: E14 soma g1 and E14 soma g2 (Fig.1B and Fig. S1B). In E14 Soma g1 cells, we observed high expression of primitive streak (PS) and embryonic mesoderm genes: *T, EOMES* and *MESP1*, the cell surface markers: *KDR, PDGFRA, CXCR4* and CD13 (*ANPEP*) (Kopper and Benvenisty, 2012), and the signalling components: *WNT5A, WNT8A* and *LEF1*. These cells also showed high levels of *SNAI1, ZEB2* and *CDH2* (N-Cadherin)(Pan et al., 2016) and low expression of *CDH1* (E-cadherin). The gene expression profile in soma g1 suggests these cells may be undergoing epithelial– mesenchymal transition (Stemmler et al., 2019). In contrast, soma g2 cells in E14 embryos exhibit epithelial features, with hallmark expression of amnion specific genes *GATA3, GATA2, TFAP2A, TFAP2C, OVOL1, KRT7/8/1*8 (Gomes Fernandes et al., 2018; Xiang et al., 2020), as well as cell adhesion-related genes *ITGA3, PKP2, PODXL* and *AHNAK* (Saykali et al., 2019). Trajectory analysis confirmed the pseudo-temporal relationship amongst these cells, with soma g1 nascent mesoderm being closer to epiblast cells, while soma g2 diverge from g1 and PGCs (Fig. S1C). There is evidently a close spatial relationship between pre-migratory pPGCs, mesoderm and amnion precursors (see later). Previous studies have shown that after their induction in the posterior early-PS epiblast, the PGC cluster localizes at the embryonic and extraembryonic border in pig pre-somitic stage embryos (Kobayashi et al., 2017; Wolf et al., 2011). Similarly, in a gastrulating CS7 human embryo, hPGCs are suggested to emerge from PS and are set aside from nascent mesoderm and other lineages (Tyser et al., 2020). Importantly, all these cell types are induced by BMP signalling, which is detected in the posterior end of the pig embryo from E12 onwards (Valdez Magana et al., 2014; Yoshida et al., 2016).

Next, we identified differentially expressed genes (DEG) between E14 PGC and E14 soma (g1 and g2 combined) and found enrichment in PGCs for “germ cell development” and “positive regulation of double-strand break repair” by gene ontology (GO) analysis (Fig. S1D and Table S2), indicating the importance of DNA repair during pre-meiotic PGC development (Guo et al., 2015; Hajkova et al., 2010; Hill and Crossan, 2019). Furthermore, GO analysis showed reduced expression of glycolysis-associated genes in PGCs between E14 and E31, and upregulation of genes controlling mitochondrial activity and oxidative phosphorylation (Fig. S1E, Fig. S2A and Table S2). An increase in mitochondrial activity in E31 gonadal PGCs is also suggested by higher expression of mtDNA-encoded genes compared to E14 (Fig. S1E). Thus, these results are consistent with a metabolic shift in pPGCs during their migration and epigenetic resetting as reported previously in mouse (Hayashi et al., 2017) and human gonadal PGCs (Floros et al., 2018). Notably, these metabolic changes start in pre-migratory pPGCs, supporting previous observations in hPGCLCs (Tischler et al., 2019).

To gain insight into the signalling microenvironment of the posterior end of E14 pig embryos, we analysed the expression profile of genes involved in different signalling pathways. GO terms and KEGG pathways analysis of E14 somatic compartment showed enrichment for WNT, BMP, TGFβ, and PI3K-akt signalling (Fig. S1D and Fig. S2B), similar to findings from pre-streak and early PS pig embryos (E10.5-12.5)(Valdez Magana et al., 2014; Yoshida et

al., 2016). Previous work showed that WNT signalling confers pig germ cell precursors the competence to respond to BMP and triggers the germ cell program at around E12 (Kobayashi et al., 2017; Kojima et al., 2017). We show that after the onset of pPGC specification these key signalling molecules are still expressed in this area of development of the extraembryonic mesoderm, which gives rise to amnion (Perry, 1981).

In contrast to the soma, from the earliest developmental stage (E14) pPGCs showed upregulation of Jak/STAT-Insulin pathways genes (Fig. S2B), which is consistent with the described function of LIF as a survival factor in PGCs (Hayashi et al., 2011; Ohinata et al., 2009).

We next examined the cell cycle stage of pre-migratory pPGCs and determined that more than 85% of cells were in either G1 or G2 cell cycle stage, in contrast to their early gonadal counterparts that were mostly in S-phase (46%) (Fig. S2C). These findings are in line with previous observations that E14 pPGCs cease to incorporate EdU and that a high proportion of migratory E17 pPGCs are arrested in G2 phase of the cell cycle, suggesting that pre-gonadal PGCs do not proliferate rapidly (Hyldig et al., 2011a; Kobayashi et al., 2017). This kinetics are also consistent with limited proliferation of hPGCLCs during the first days (day 4) of development, which then resumes during extended culture (Gell et al., 2020).

To investigate the conservation of germline development in detail we compared the expression profiles of pPGCs, hPGCs and *Cynomolgus* monkey PGCs (cyPGCs) by integrating scRNASeq datasets (Li et al., 2017; Sasaki et al., 2016). E14 pPGCs clustered mostly with Wk4-5 hPGCs and E13-20 cyPGCs (ePGCs), whereas E31 pPGCs clustered with Wk5-7 hPGCs and E36-55 cyPGCs (lPGCs) (Fig. 1D and Fig. S1F). Human, pig and monkey PGCs show similar expression profiles for key germline genes (*SOX17, PRDM1 (BLIMP1), TFAP2C, NANOS3, DND1*) and pluripotency genes (*NANOG, POU5F1, SOX2, ESRRB, LIN28A and ZIC3*) (Fig. 1E; Table S3). As in hPGCs and cyPGCs, the endoderm marker *GATA4* is also widely expressed in pPGCs, the mesoderm marker *T (BRACHYURY)* is expressed in early pPGCs and maintained in some gonadal pPGCs, whereas *EOMES* is absent from pre-migratory cells (Fig. 1B). Conversely, the naïve pluripotency gene *TFCP2L1* is not detectable and *KLF4* is only found in few pPGCs (Fig. 1E). Although expression of these genes occurs in the human and *Cynomolgus* PGC lineage, both seem to be dispensable for hPGCLC specification (Hancock et al., 2020). Similarly, *PRDM14* is only detectable in gonadal pPGCs, and may not have an essential role during pPGC specification (Fig. 1B, Fig. 1E) (Kobayashi et al., 2017). Recent evidence shows that *PRDM14* supports hPGC number rather than specification in humans (Sybirna et al., 2020).

This analysis shows that the transcriptional program for pPGC specification is largely equivalent to that of hPGCs and cyPGCs, but differs from that of mice (Fig. 1F) (Guo et al., 2015; Irie et al., 2015; Kojima et al., 2017; Sasaki et al., 2016). Although the basis for the transcriptional divergence is not fully understood, it is noteworthy that pigs and humans (and most other mammals) develop a bilaminar disc prior to the onset of gastrulation, whereas some rodents, like mice and rats, have evolved an egg cylinder. The divergence in development and molecular aspects such as the pluripotency network, which may facilitate the evolution of embryological innovations, merits further consideration (Johnson and Alberio, 2015).

The reduced (*KLF4, PRDM14*) or lack of expression (*SOX2, TFCP2L1*) of some of these genes in the pig germline prompted us to investigate underlying pluripotency features of pPGCs in more detail. We created signature gene sets from E6 ICM, as well as E8 and E11 epiblast (Ramos-Ibeas et al., 2019) and examined their expression in pre-migratory (E14) and gonadal pPGCs (E31). A strong pig E8 Epi signature score was determined for both (E14 and E31) pPGC stages, with gonadal pPGCs showing a stronger ICM signature score compared to E14 pPGCs (Fig. 1G and Table S4). The signature genes contributing to these scores include elevated expression of well-known transcription factors (*POU5F1, NR5A2, SOX15*), but also of chromatin related genes (*HELLS, BRDT, ZAR1*) and regulators of transposable elements activity (*MOV10, ASZ1, PLD6, HENMT, TDRKH* and *SAMHD1*), indicating that restoration of a gene signature common with ICM/E8 Epi in the early PGCs is linked to epigenetic resetting of the germline, which does not occur in the neighbouring somatic lineages.

Membrane proteins participate in numerous cellular processes, such as cell signalling, transport and migration. Therefore, we sought to identify pPGC-specific membrane proteins by selecting pPGC-specific genes with relevant GO terms and/or curated in the Cell Surface Protein Atlas (Bausch-Fluck et al., 2015). As reported before in hPGCs and early cyPGCs (Gomes Fernandes et al., 2018; Sasaki et al., 2016; Tang et al., 2015), *KIT* and *PDPN* were upregulated in pre-migratory pPGCs (Fig. S2D). We also determined expression of the orphan receptor *GPR50* which is specific for early but not gonadal pPGC (Fig. S2E). By IF, GPR50 was detected on the cell membrane of early migratory pPGCs, however gonadal stages show strong nucleolar staining (Fig. S2F). GPR50 is an orphan receptor shown to promote cell migration (Wojciech et al., 2018). We also detected high levels of *CXCR4*, needed for PGC migration in mice (Molyneaux et al., 2003), in E14 pPGCs suggesting the onset of migration (Fig. 1B). GDF3, a mammalian-specific TGFβ ligand expressed in *cy*PGCs (Sasaki et al., 2016) and gonadal hPGCs (Li et al., 2017), is also enriched in early pPGCs (Fig. S2D). CD markers including *CD126* (*IL-6R*), *CD157 (BST1)*, closely related to the hPGC marker *CD38*, and the orphan receptor *GPR133* (*ADGRD1)*, which is also expressed in hPGCs, are all upregulated in pPGC (Fig. S2D, S2E). Additionally, upregulation of *SLC23A2* in pre-migratory PGC may contribute to the cellular uptake of vitamin C and promote TET1 activity in PGCs (DiTroia et al., 2019). Altogether, the surface molecules identified depict a profile of cells preparing to embark on their migration towards the gonad, and onset of epigenetic resetting.

### Onset of DNA demethylation in Pre-migratory pPGCs

Next, we investigated the onset of epigenetic reprogramming in pPGC using a combination of approaches. Analysis by IF showed 5-hydroxymethylcytosine (5hmC) staining in E14 pPGCs, concomitantly with reduced 5-methylcytosine (5mC) (Kobayashi et al., 2017), suggesting the onset of DNA demethylation (Fig. 2A). Quantification of 5mC and 5hmC, using liquid chromatography–tandem mass spectrometry (LC-MS/MS) (Hill et al., 2018), was consistent with the IF data where we demonstrated that 5hmC levels are higher in pre-migratory (E14) pPGC compared to the surrounding cells and Epi. Conversely 5mC levels are lower in pre-migratory pPGC compared to Epi (Fig. 2B). DNA methylation reaches the lowest levels in gonadal pPGCs (Fig. 2B and Fig. S3B). Importantly, we also determined a similar kinetic of 5mC and 5hmC in D4 hPGCLCs and equivalent human gonadal samples (Fig. 2B), in accordance with previous reports in early gonadal hPGCs (Guo et al., 2015; Tang et al., 2015). Coupled with the high levels of 5hmC, we detected a sharp decline in *DNMT3B* and *UHRF1*, indicating that the methylation machinery is downregulated from the pre-migratory stage and persists until gonadal stages (Fig. 2C and Fig. S3A).

**Figure 2.**
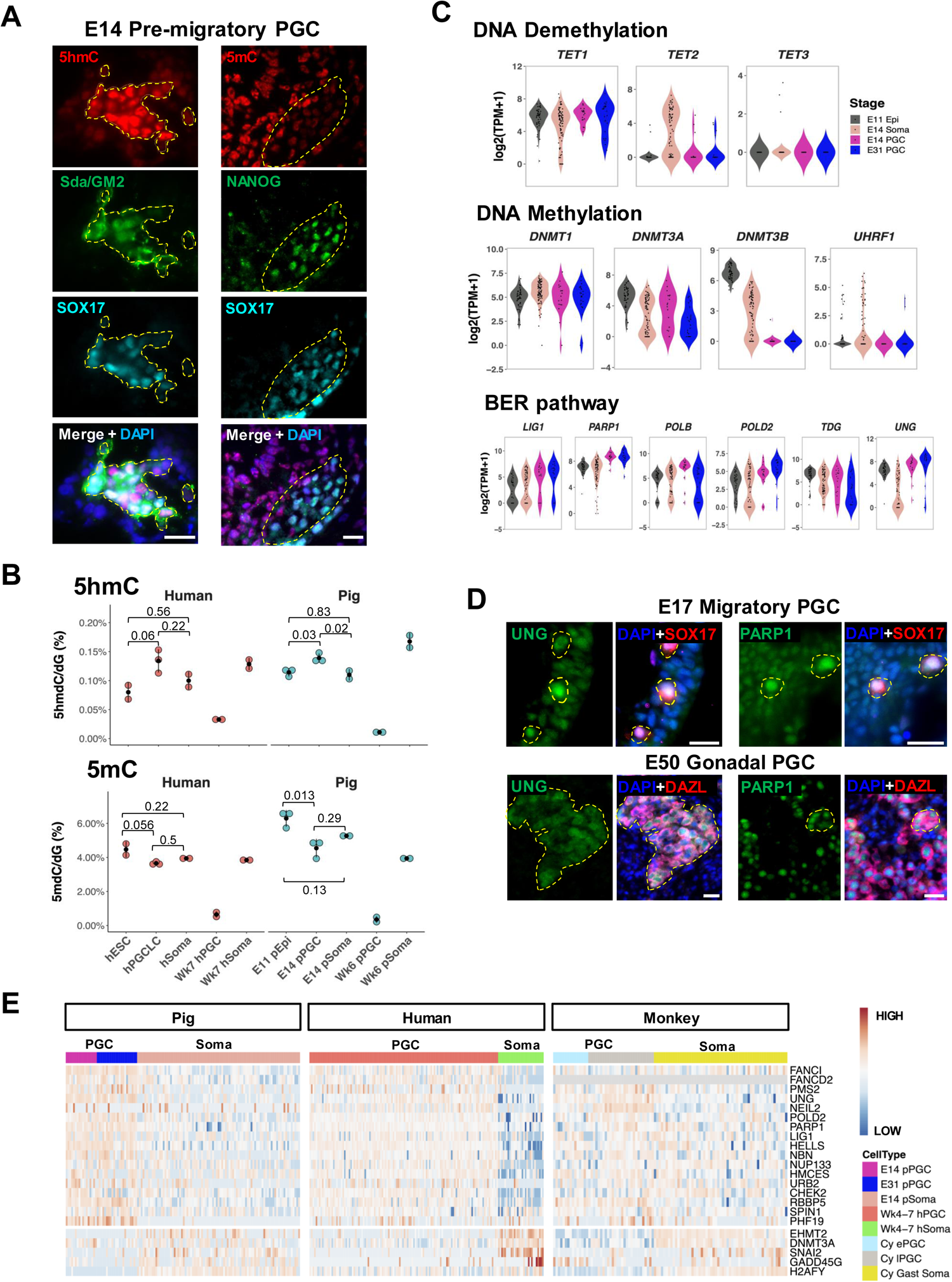
Active DNA Demethylation in Pre-migratory pPGC. (A) Immunofluorescence of 5hmC and 5mC in E14 PGC cluster (indicated by yellow dashed lines). PGCs are marked by SOX17 and Sda/GM2 and Nanog. Scale bar: 20µm. (B) 5hmC and 5mC levels determined by LC–MS/MS. Methylation levels are indicated relative to total levels of deoxyguanine (dG). *P* values are based on combined ANOVA and Holm’s post hoc test. (C) Expression of epigenetic modifiers for DNA methylation/demethylation and BER pathway components in E11 Epiblast, E14 Somatic Cells, E14 PGC and E31 PGC. (D) Immunofluorescence of BER pathway component UNG (left) and PARP1 (right) in E17 migratory and E50 Gonadal PGC. Yellow circle indicates PGC. Scale bar: 20µm. (E) Expression heatmap of epigenetic modifiers that show differential expression in pPGC, hPGC and cyPGC compared to somatic tissue. “Cy ePGC”: early cyPGC (E13-20); “Cy lPGC”: late cyPGC (E36-55); “Cy Gast Soma”: cy gastrulating cells (E13-20). Grey colour in heatmap indicates “NA”. As different expression units are used for the three species, values in the colour scale are replaced by HIGH and LOW. See also Figure S3.

We also found that multiple base excision repair (BER) pathway genes (*LIG1, POLD2, POLB, PARP1 and UNG)*, were upregulated in E14 and E31 pPGCs (Fig. 2C and D), supporting the suggestion that the active removal of TET-oxidized products in PGC may be mediated by the BER pathway (Hackett et al., 2013; Hajkova et al., 2010; Hill et al., 2018). Furthermore, upregulation of ‘readers’ for TET-oxidized products (*HELLS, HMCES, NUP133 and URB2*) (Spruijt et al., 2013) was observed in pPGCs, cyPGCs and hPGCs, suggesting that 5hmC may be a dynamic and functional marker in early PGCs (Fig. 2E). In addition to the BER pathway, we detected upregulation of components of Fanconi Anaemia (FA) (*FANCI and FANCD2)*, mismatch repair *(PMS2)* and double-strand break repair *(NBN)* pathways in pPGC, indicating that multiple DNA repair mechanisms may be activated during epigenetic reprogramming of pre-migratory PGCs (Fig. 2C and S2E). Altogether, our data from IF, LC-MS/MS, and scRNASeq show that non-replicative pre-migratory pPGCs initiate TET activities and activate BER pathway components potentially mediating active DNA demethylation, followed by passive demethylation in migratory and gonadal PGCs, as shown by the reduction in DNMT3A/B and UHRF1. This indicates that DNA demethylation is controlled by active and passive mechanisms that start in early PGCs (E14), which reach the lowest levels in gonadal stages. These mechanisms cannot be studied in human nascent PGCs, however our findings in the pig concur with those reported previously showing limited DNA replication (Gell et al., 2020) and high levels of 5hmC in D4 hPGCLCs cells (Tang et al., 2015), and increased expression of BER pathway genes in Wk4 hPGCs (Guo et al., 2015).

The extended DNA demethylation kinetic in the pig (∼21 days) contrasts with the rapid demethylation in mouse PGCs (∼5 days), where it is primarily mediated by passive demethylation during early migration, followed by active and passive demethylation in the gonads (Hackett et al., 2013; Hill et al., 2018; Kagiwada et al., 2013). The protracted process in the pig germline reflects the longer period of development of pig and human PGCs, which are specified around Wk2 and reach the gonadal ridges at Wk4 and Wk5, respectively (Takagi et al., 1997; Witchi, 1948); in the mouse this process takes ∼4 days (from E6.25-10.5). Yet, the number of PGCs in the early gonad is similar between species (∼2600 in mouse E11.5 (Kagiwada et al., 2013), ∼ 3000 in Wk5 human male fetal gonad (Bendsen et al., 2003), ∼3-5000 in pig Wk4 gonad (Black and Erikson, 1965; our unpublished data). To reach the same number of gonadal germ cells, mouse PGCs proliferate faster and divide every ∼12 hrs, whereas human PGCs divide every 6 days (Bendsen et al., 2006; Kagiwada et al., 2013). Thus, in the context of prolonged doubling times in humans and pig PGCs, complementary DNA demethylation mechanisms (active and passive) apparently ensure efficient initiation of DNA methylation reprogramming.

### Dynamic chromatin changes in pPGC

We next examined chromatin features of pPGCs, as part of the epigenetic resetting and DNA demethylation in pPGCs. Whilst overall H3K27me3 was elevated in migratory (E17) and early gonadal (E25) PGCs, it decreased sharply in mid- and late gonadal PGCs (Fig. 3A and 4A), consistent with the high expression of *Polycomb related complex 2* (PRC2) members *EZH2, SUZ12* and *EED* in migratory and early gonadal pPGC (Supp. Fig. 3C). Furthermore, the PRC2 associated cofactor *PHF19*, required for PRC2 recruitment and activity (Ballaré et al., 2012), was enriched in early pPGC (Fig. S3C). Changes in other histone and chromatin remodellers were also detected, such as the upregulation of components of MII complex (*DPY30, RBBP5*) and SWI/SNF proteins *SMARCA5* and *HLTF* (Fig. S3C). Similar observations have been reported in Wk4 hPGCs and D4 hPGCLCs (Gell et al., 2020; Gkountela et al., 2013; Gomes Fernandes et al., 2018; Tang et al., 2015). By contrast, mouse PGCs show persistent H3K27me3 in gonadal PGCs (de Sousa Lopes et al., 2008; Seki et al., 2005), which might have a role in maintaining genomic integrity during the period of active DNA demethylation (Cotton et al., 2013). The decreases in H3K27me3 in gonadal PGCs in the pig and human during extensive DNA demethylation suggests possible additional mechanisms that warrant future investigations.

**Figure 3.**
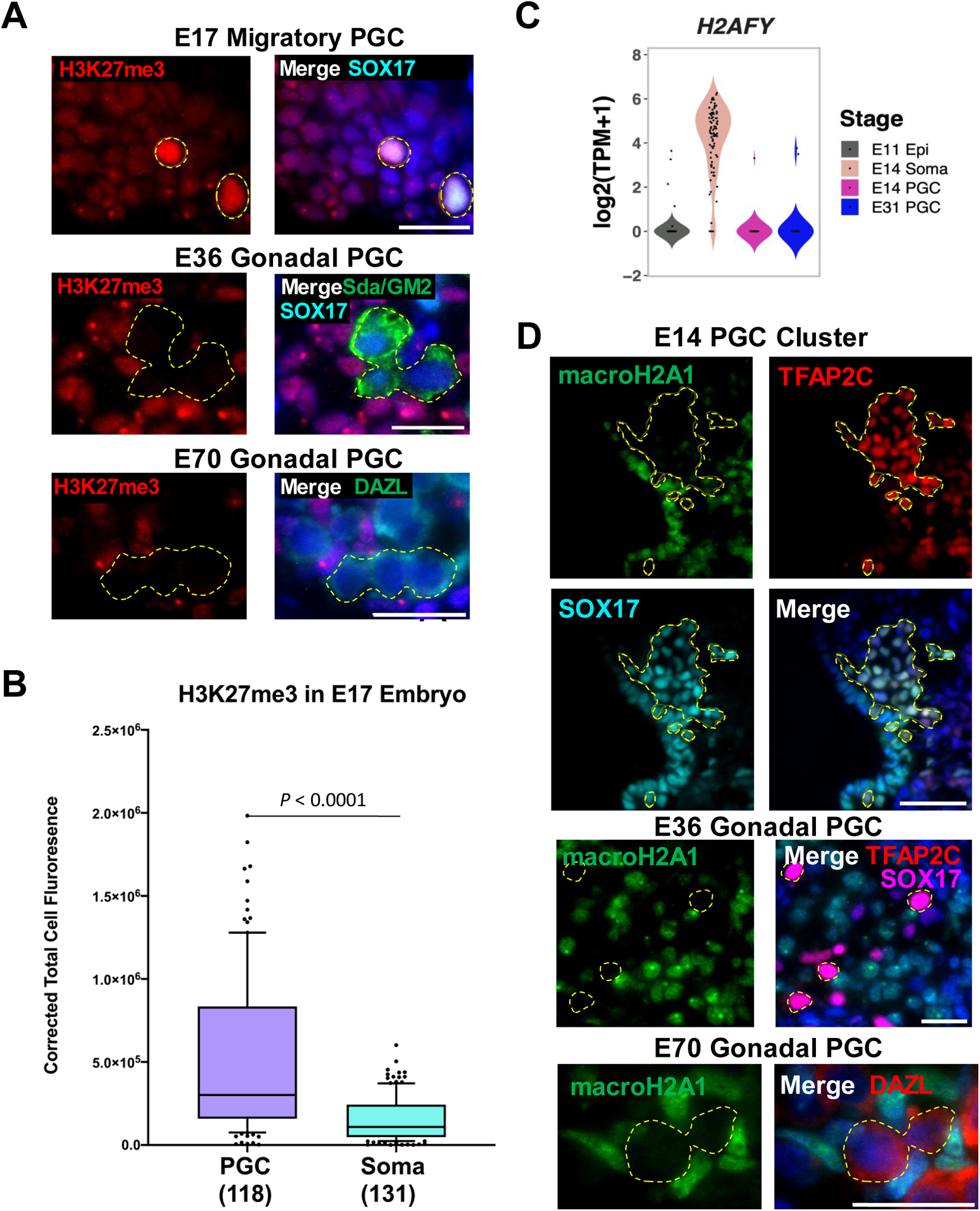
Histone remodelling in Pre-migratory, early migratory and gonadal pPGC. (A) Immunofluorescence of H3K27me3 in E17 migratory, E36 and E70 gonadal PGC. Yellow circle indicates PGC. PGCs are marked by SOX17 and/or Sda/GM2 in E17 and E36, and DAZL in E70. Scale bar: 20µm. (B) Quantification of H3K27me3 in E17 migratory PGC and surrounding somatic cells (boxes, mean and interquartile ranges; whiskers, 10-90 percentile; Mann–Whitney *U*-test). (C) Expression profile of *H2AFY*, which encode macroH2A1 in E11 Epiblast cells, E14 Somatic cells, E14 PGC and E31 PGC. (D) Immunofluorescence of macroH2A1 (*H2AFY*) in E14 PGC cluster, E36 and E70 Gonadal PGCs. PGCs are marked by SOX17 and TFAP2C in E14 and E36, and DAZL in E70. Scale bar: 20µm. Yellow circles indicate PGC. See also Figure S3.

MacroH2A1, the macro-histone variant encoded by *H2AFY* associated with H3K27me3 on developmental genes, was upregulated in somatic cells, but not in pPGCs, where it would act as a barrier to transcription factor-induced reprogramming (Gaspar-Maia et al., 2013) (Fig. 3C, 3D). MacroH2A1.1 modulates PARP1 activity and mediates cellular DNA damage response (Posavec Marjanović et al., 2017). Interestingly, we found high PARP1 levels in pre-migratory and gonadal PGCs (Fig. 2C-E), suggesting that macroH2A depletion from early PGCs might contribute to the maintenance of a chromatin configuration that facilitates the onset of epigenetic reprogramming. Consistent with the findings in pPGCs, *H2AFY* is downregulated in human and cyPGCs (Fig 2E). Furthermore, gonadal hPGCs were shown to lack the closely related macroH2A2 (Tang et al., 2015).

### Extensive X chromosome reactivation in pre-migratory pPGC

To gain further insight into reprogramming in pre-migratory pPGCs we combined IF and transcriptomic analysis of XCR, which is characterized by the loss of H3K27me3 enrichment on the Xi and bi-allelic expression of X-linked genes (Sugimoto and Abe, 2007). We found that in pre-/early migratory (E14-17) and gonadal pPGCs (E25) over 70% of cells showed faint or no H3K27me3 “spots” (Fig. 4A, Fig. S4C and S4D), suggesting XCR had already started in female pre-migratory cells. Notably, the histone demethylase *KDM6A*, which is associated with the loss of H3K27me3 in the inactive X chromosome (XC) (Borensztein et al., 2017; Mansour et al., 2012), was upregulated in E14 female PGCs (Fig. 4B). To further analyse XCR at the transcriptional level, we measured *XIST* expression, which is critical for X inactivation (Jonkers et al., 2008). After determination of the sexual identity of E14 and E31 cells based on the cumulative levels of Y chromosome genes per cell (Supp. Fig. 4A), we determined a reduction in *XIST* expression in the majority of E14 (4/6) and E31 (5/8) female pPGCs, but not in female somatic cells (Fig. 4C). Furthermore, XC, but not autosome, expression in female E14 PGCs was significantly higher compared to male pPGCs, increasing further in E31 female PGCs (Fig. 4D). In contrast, no gender differences were detected for either XC or autosome expression in somatic cells (Fig. 4D). At the single cell level, the XC expression to total autosomal expression (X:allA) ratio was above 1 in all E31 female PGCs and most E14 female PGCs (Fig. S4B), consistent with observations in female gonadal mPGC and hPGC (Sangrithi et al., 2017). Also, we found no apparent relationship between X-linked gene reactivation and the proximity to the XC inactivation (XCI) centre (Fig. 4E)

**Figure 4.**
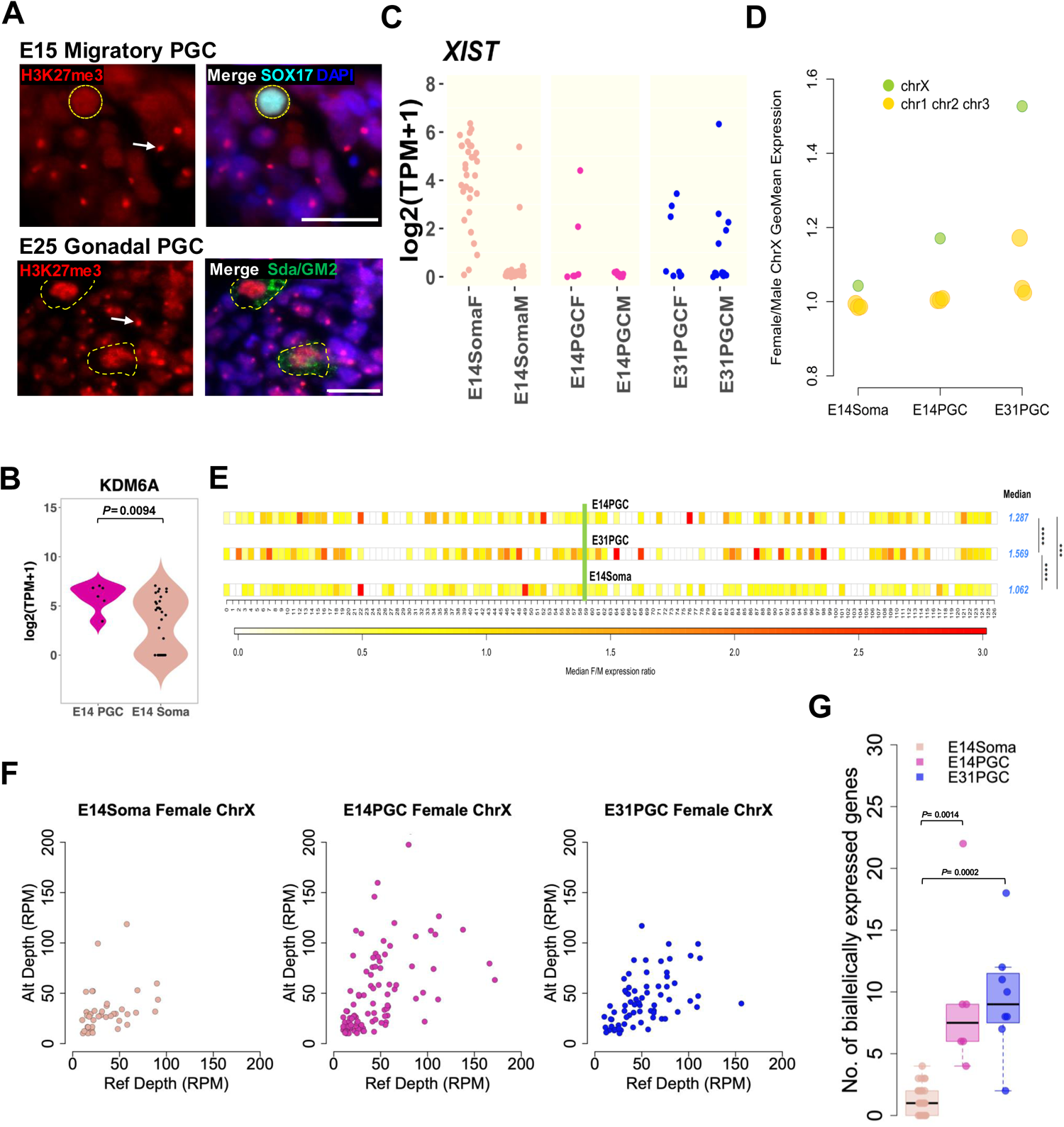
X chromosome Reactivation in Pre-migratory pPGC. (A) Immunofluorescence of H3K27me3 in E15 early migratory and E25 early gonadal pPGC. Xi-associated H3K27me3 is detected in somatic cells (arrows). Yellow dashed circle marks PGC. Scale bar: 20µm. (B) Expression of KDM6A in E14 cells. *P* value is determined by Mann-Whitney U-test. (C) Expression of XIST in E14 Somatic Cells, E14 PGCs and E31 PGCs. (D) Female to male expression ratio of X chromosome genes vs. autosomes (chr1, chr2 and chr3) in E14 somatic cells, E14 PGCs and E31 PGCs. (E) Median female to male expression ratio across X chromosome in E14 PGC, E31 PGC and E14 Somatic cells. p value. *** p < 0.001; **** p < 0.0001 by pairwise Wilcoxon test. Green line indicates estimated position of X chromosome inactivation centre. (F) Biallelically-detected SNPs on X chromosome identified in female E14 somatic cells, E14 PGCs, and E31 PGCs. Each dot represents one biallelically-detected SNP. X axis: Sum of reads (RPM) that are mapped to the reference alleles; Y axis: Sum of reads (RPM) that are mapped to the alternative alleles. (G) Number of biallelically expressed genes in E14 PGCs, E31 PGCs and E14 Somatic cells. *P* value is determined by Kruskal-Wallis test followed by Dunn’s test. See also Figure S4.

To rule out the possibility that the increased X:allA ratio and F:M ratio for XC was due to expression changes in one active XC, instead of biallelic expression from both XCs, we analysed gene expression at allelic resolution. E14 female somatic cells have a lower number of bi-allelic single nucleotide polymorphisms (SNPs) (Fig. 4F), which are likely to be genes that escape XCI in pig. Studies show that 4-8% and 15-25% of X-linked genes in mice and in human, respectively, escape XCI to some degree (Carrel and Willard, 2005). These genes, which we called XC “*escapers”* to distinguish them from the DNA methylation escapees (see later), vary largely between tissues and species and have not been characterized in the pig. Therefore, we categorized X-linked genes containing biallelic SNPs in pig somatic cells as our XC *escapers*. Consistent with the increased X:A and F:M ratio, E14 and E31 female PGCs have a large number of non-escaper, biallelic SNPs, providing evidence of onset of XCR in pre-migratory PGCs (Fig. 4F). We then identified biallelically expressed X-linked genes in female cells and found that all female pPGCs contain at least one biallelically expressed X-linked gene that is not found in somatic cells. In contrast to the sharp increase of biallelic gene expression, which is only detected in mouse gonadal PGCs (Sugimoto and Abe, 2007), both pig pre-migratory and gonadal PGCs have higher number of biallelically expressed genes, suggesting that XCR is a cell-autonomous and asynchronous process taking place over a long period (Fig. 4G).

Consistent with our findings in pPGCs, hallmarks of XCR have also been reported in hPGCs, showing loss of the H3K27me3 spot at Wk4 (Tang et al., 2015), and biallelic expression of X-linked genes at Wk7-8 PGCs; however data from earlier stages is not available (Sangrithi et al., 2017; Vértesy et al., 2018). Even though it is not currently possible to conclude whether human XCR occurs as early as shown in pPGCs, our evidence of XCR in pig pre-migratory pPGCs contrasts with observations in mouse PGCs, where there is limited loss of H3K27me3 (<10%) and *Xist* (<15%) expression in pre-migratory PGCs; the increase in X:A ratio is first detected in E11.5 PGCs (de Sousa Lopes et al., 2008; Sangrithi et al., 2017; Sugimoto and Abe, 2007). Taken together, our findings show that XCR begins in pre-/early migratory pPGCs and continues in gonadal pPGCs.

### DNA methylation level reaches basal level in gonadal pPGC

We sought to obtain detailed information on pPGC demethylation by generating whole genome base-resolution PBAT libraries of Wk5 (E35) gonadal pig PGC from 2 female and 2 male embryos (Table S1). In each replicate, over 90% of total genomic CpG sites were detected (i.e. covered with at least one read) and nearly 60% (apart from one sample of somatic cells (Soma.female), which is 52%) were covered by at least five reads (5x). The bisulfite conversion rate was around 99%, determined with the spiked unmethylated lambda DNA (Table S5). Consistent with the LC-MS results (Fig. 2B), Wk5 pPGCs reach basal levels of DNA methylation (around 1%) in both genders, whereas equivalent gonadal somatic cells showed a median level of over 75% methylation (Fig. 5A). Extensive DNA demethylation was determined across all genomic features, including CpG islands (CGI), promoters, introns, intergenic regions and exons (Fig. 5B and 5C). Furthermore, Wk5 PGCs also showed comprehensive demethylation of imprinted genes (Fig. 5D), except for *PEG10*, which retained some methylation (7-15%). The loss of DNA methylation at most imprinted loci in early gonadal germ cells is in line with previous reports showing that DNA demethylation at imprinted loci starts prior to the arrival to the genital ridges (Hyldig et al., 2011a; Petkov et al., 2009).

**Figure 5.**
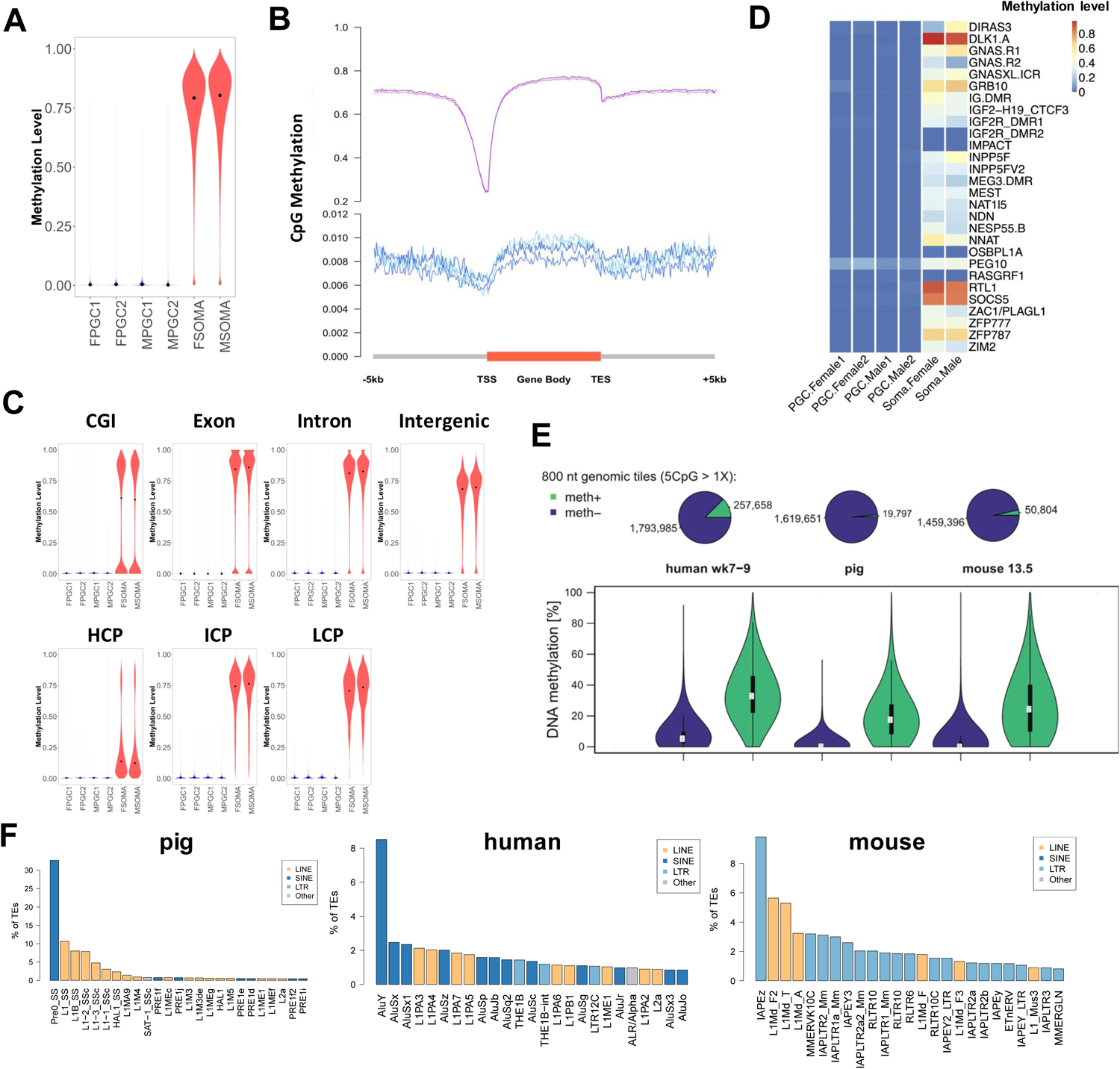
PBAT reveal basal level of methylation in gonadal pPGCs. (A) CpG methylation levels in 1 kb genomic tiles of Wk5 (E35) female and male pPGCs and gonadal somatic cells. Black point indicates median. (B) Averaged CpG methylation level profiles of all genes from 5 kb upstream (-) of transcription start sites (TSSs), through scaled gene bodies to 5 kb downstream (+) of transcription end sites (TES). Different y axes are used for pPGCs and somatic cells due to extreme low level of methylation in pPGCs. (C) Violin plots showing CpG methylation levels in different genomic features in samples indicated. (D) CpG methylation levels of imprinted regions in pPGCs and somatic samples. (E) Top: Proportion of demethylated loci (meth-) and demethylation-resistant loci (meth+) in Wk5 pPGCs, Wk7-9 hPGCs and E13.5 mPGCs (the number of meth+ and meth-800nt genomic tiles are indicated on the pie chart). Bottom: CpG methylation levels of meth- and escapees (meth+) in three species. White dot indicates median, black bar indicates interquartile range. (F) Distribution of TE families that overlap with TE-rich escapees in Wk5 pPGCs, Wk7-9 hPGCs and E13.5 mPGCs. See also Figure S5 and Table S5.

Analysis of transposable elements (TE), which are extensively demethylated in mouse and human gonadal PGCs (Hajkova et al., 2002; Seisenberger et al., 2012; Tang et al., 2015), also showed very low levels of DNA methylation in male and female pPGC (Fig. S5A), consistent with previous locus specific analysis (Hyldig et al., 2011a; Petkov et al., 2009). DNA demethylation was concurrent with increased expression of major TE families, including long and short interspersed elements (LINEs and SINEs) and long terminal repeats (LTRs) in E14 and E31 (Wk5) pPGCs (Fig. S5B), in line with reports in human and mouse gonadal PGCs (Guo et al., 2015; Ohno et al., 2013). A concomitant up-regulation of negative regulators of TE activity in PGCs, including the piRNA pathway, suggests that the mobilization of retrotransposons is likely to be repressed despite of the increased expression of TE elements (Fig. S2A and Table S6).

The overall low-level DNA methylation in Wk5 pPGC (∼1%) was comparable with that of Wk7-9 hPGC (∼4.5%) and E13.5 mPGC (2.5%) (Kobayashi et al., 2013; Tang et al., 2015). Despite the comprehensive demethylation, a small proportion of loci still maintain partial methylation (Fig. 5E), as in mouse and human (Guibert et al., 2012; Seisenberger et al., 2012; Tang et al., 2015). A large proportion of these loci are found located within TE rabundant regions, whose distribution in the genome is variable, thus influencing the overall methylation levels in each species (Fig. S5C and S5D). In the pig, the relative content of TE (∼40%) in the genome is lower than that of other mammals (Fang et al., 2012; Groenen et al., 2012), which could explain the reduced number of demethylation-resistant loci identified in this species (Fig. 5E). We identified high-confidence demethylation-resistant loci as “escapees”. Notably, the most abundant repeat families at TE-rich (>=10% overlap with TE) escapees are species-specific and evolutionarily young TE, including the pig SINE element Pre0_SS of the PRE1-family, human AluY, and mouse IAPEz repeats (Fig. 5F). The overall observations in pig germ cells on global DNA demethylation and resistant loci parallel those in human and mice.

### TE-poor escapees show overlapping features between species

Many pig escapees at TE-poor (<10% overlap with TE) regions are associated with promoters, CGI and gene bodies, as in human and mouse PGCs (Kobayashi et al., 2013; Tang et al., 2015). Their numbers vary, with the lowest in mPGC (1,059), compared to pPGC (1,402) and hPGC (6,009) (Fig. 6A). The larger proportion of TE-poor escapees (13%, 1,402/10,421) in pPGC could be due to the relatively lower content of repetitive elements in the pig genome (Fang et al., 2012; Groenen et al., 2012) (Fig. 6A). Nearly 21.5% (44/205) of TE-poor escapee regions in pig show conserved synteny with human, compared to 4% (8/206) in mouse (Fig. 6B). In addition, we found that 265 (47%) TE-poor escapee genes in pig and 191 (23.2%) in mouse are in common with human escapee genes (Fig. 6C) (Tang et al., 2015). Comparison with the NHGRI GWAS catalogue revealed that the 265 human-pig conserved TE-poor escapee genes are linked with metabolic and neurological traits such as obesity-linked disorders and schizophrenia (Fig. S6A). Some of the disease-associated genes show sequence conservation between human and pig, such as obesity-related gene *SORCS2* and schizophrenia-related *PLCH2* (Fig. 6D). For pig specific TE-poor escapee genes, the comparison with the GWAS catalogue revealed pig specific terms, such as association with asthma (Fig. S6B). TE-rich escapee genes overlapping with pig-specific TEs (*Pre0_SS* and *L1_SS*) also show enrichment for developmental-, metabolic- and neurological-related GO terms, such as *FTO*, an obesity-related gene (Fig. 6D and Fig. S6C).

**Figure 6.**
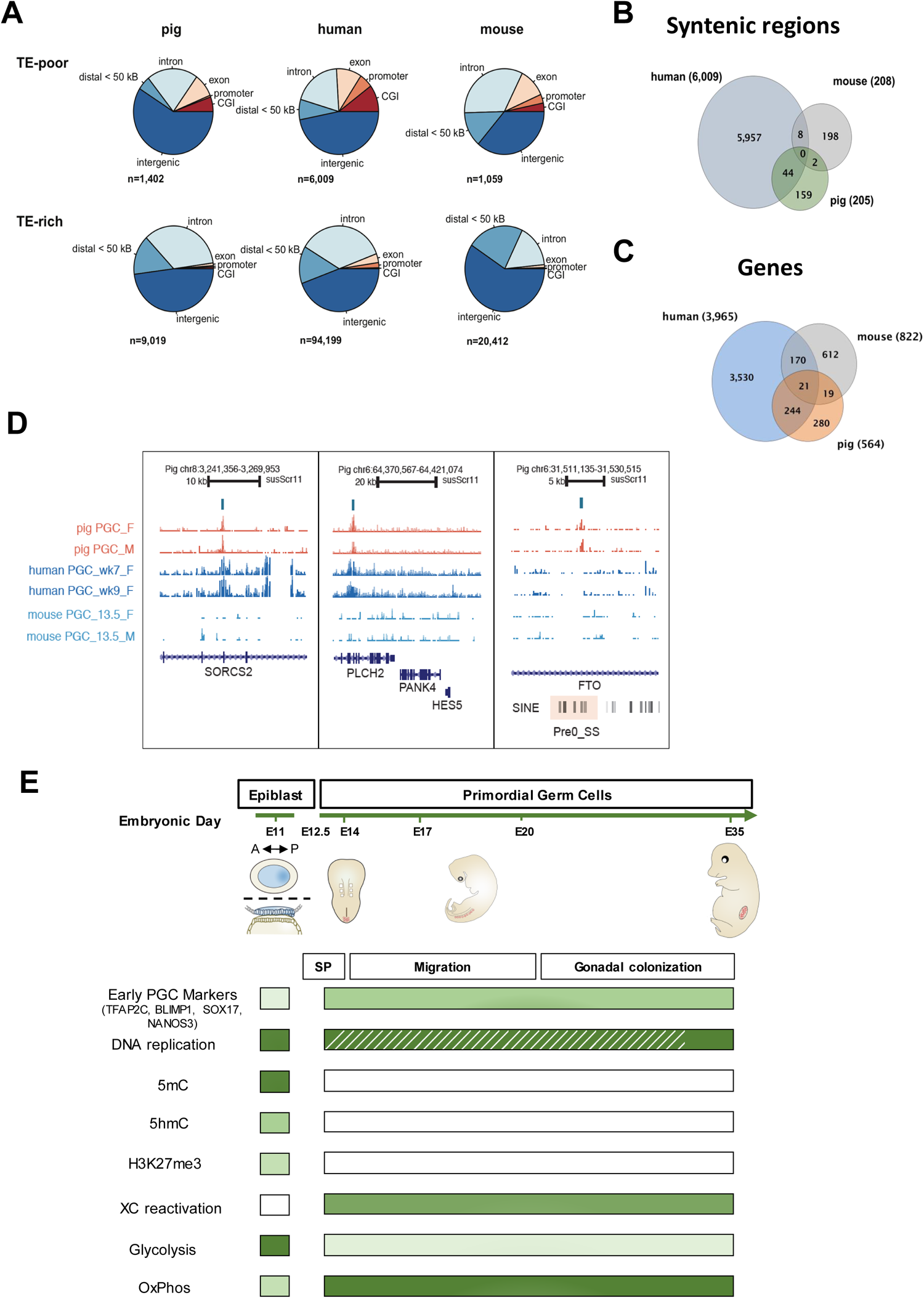
Common and unique features in DNA demethylation escapees between mouse, human and pig. (A) Distribution of TE-poor (<10% overlap with TE) and TE-rich (≥10% overlap) escapees in Wk5 pPGCs, Wk7-9 hPGCs and E13.5 mPGCs. Number of escapees (n) were identified by methylation level (at least 30% in human and 15% in pig and mouse). (B) Overlap of syntenic TE-poor escapees among pig, human and mouse. Escapee regions in pig (205) and mouse (208) were lifted over to compare with syntenic regions in the human genome. (C) Overlap of homologous, TE-poor escapee genes among pig, human and mouse. (D) The TE-poor escapee regions within SORCS2 and PLCH2 are conserved between human and pig, while a pig-specific escapee is identified within FTO. (E) Diagrammatic representation of the events in the pig germline. SP: indicates germline specification; OxPhos: indicates oxidative phosphorylation; The dash line indicates expected DNA synthesis in migratory and early gonadal PGCs. See also Supplementary Figure 6.

Lastly, analysis of common TE-poor escapee genes across at least two species (pig, human, and mouse) revealed enrichment for brain-specific gene expression, consistent with their association with neurological-related traits. These common genes also showed enrichment for key protein domains in the KRAB-ZFP family, suggesting a conserved mechanism for maintenance of methylation at these loci across species (Fig. S6D).

## Conclusions

Our investigation advances insight into the mechanism of pPGCs specification, and their subsequent development. Notably, pPGC specification is closely linked with the initiation of the epigenetic program in the absence of DNA replication, a unique germline property not seen in the neighbouring somatic cells (Fig. 6E). There is likely contribution through active mechanisms of DNA demethylation as suggested by the conversion of 5mC to 5hmC, as well as the upregulation of factors of the BER mechanism. Other factors associated with DNA repair are detected in early pPGCs at the time of epigenetic reprogramming, which is crucial for the germline that transmits genetic information to subsequent generations. The erasure of 5mC would necessitate alternative host defence mechanisms for the repression of transposable elements. Passive loss of 5mC during pPGC migration is also predicated because UHRF1 is repressed in early PGCs, a crucial factor for 5mC maintenance. The detection of several cell surface markers and transcriptional changes provide a basis to unravel how the migration and subsequent development of pPGCs are regulated.

Observations on the human germline using *in vitro* models, and ex vivo hPGCs (usually after wk5) concur with the events we observe in the early pig germline. Indeed, the initial studies on the critical factors and the mechanism of hPGC specification from in vitro models, were confirmed by direct observations of pPGC specification in gastrulating pig embryos, suggesting that studies on the two species will be mutually informative. Importantly, investigations on very early human PGCs are exceptional (Tyser et al., 2020), especially during the critical period of wk2-wk4 of human development when they are essentially inaccessible. Our observations on pPGCs over this critical period covering the specification and the initiation of the epigenetic reprogramming likely apply to hPGCs.

Our study establishes a foundation for further investigations on the pig germline that will increase comprehension of the underlying developmental mechanisms. Porcine embryos are relatively accessible and ethically less challenging for studies. Genetic and other experimental approaches are possible on porcine embryos, which will lead to conceptual advances that will guide specific approaches for investigations on the human germline, including *in vitro* gametogenesis.

## Acknowledgements

Q.Z. was funded by CSC and The University of Nottingham. P.R-I was funded by Marie Sklodowska Curie Fellowship (grant agreement No 654609) and currently funded by Ramon y Cajal (RYC2018-025666-I). W.W.C.T. was supported by the Croucher Foundation. M.A.S. is supported by Wellcome Investigator Award, MRC, and core funding from Wellcome-CRUK to the Gurdon Institute. This work was supported by the Biotechnology and Biological Sciences Research Council [grant number BB/M001466/1] to R.A. and M.A.S.

## Author contributions

Q.Z, S.W., P.R-I., D.K., H.Z. performed experiments including FACS, IF, scRNASeq, embryo dissections; Q.Z and F.S. performed bioinformatics analysis. S.D. performed DNA methylation analyses; W.W.C.T. contributed with WGBS and scRNASeq protocols and analysis; C.E.R. and P.H. performed LC-MS experiments. M.L. supervised single cell analyses; R.A. supervised the project and performed dissections. R.A and M.A.S designed experiments, and together with Q.Z. wrote the paper. All authors discussed the results and contributed to the manuscript.

## Declaration of interests

The authors declare no competing interests.

## Figure Legends

**Figure S1**. Gene expression differences between pPGC and surrounding cells. Related to Figure 1.

(A) Isolation of PGCs from from E31 embryos using the cell surface marker Sda/GM2 with FACS. A section of a fetal ovary shows PGCs stained for the indicated markers (right). Scale bar: 20µm.

(B) Expression heatmap of DEGs in somatic cells found in the posterior end of E14 embryos.

(C) Expression of selected lineage markers on trajectory of single cells.

(D) Gene expression heatmap, GO terms and KEGG pathways for DEGs between E14 PGCs and somatic cells. See also Table S2.

(E) Gene expression heatmap of cellular metabolism and mitochondrial DEGs in different cells types. ETC: electron transport chain. mtDNA: mtDNA-encoded components. TCA: tricarboxylic cycle.

(F) UMAP plot showing integration of cyPGCs monkey (E13-55) (Sasaki et al., 2016) and pig (E14-31) PGCs and somatic cells.

**Figure S2**. Molecular differences between E14 and E31 pPGC, Related to Figure 1.

(A) Gene expression heatmap, GO terms and KEGG pathways for DEGs between E14 and E31 PGCs. See also Table S2.

(B) Heatmap of selected genes from different signalling pathways expressed in E14 and E31 cells.

(C) Determination of cell cycle stage based on relative average expression (module score) of canonical cell cycle markers in E14 and E31 PGCs.

(D) Expression heatmap of cell surface and membrane proteins in hPGCs, cyPGCs and pPGCs compared to somatic cells. “Cy ePGC”: early cyPGC (E13-20); “Cy lPGC”: late cyPGC (E36-55); “Cy Gast Soma”: cy gastrulating cells (E13-20). As different expression units are used for the three species, values in the colour scale are replaced by HIGH and LOW.

(E) Violin plots showing expression of cell surface proteins (CD157 (*BST1*), GPR133 (*ADGRD1*) and GPR50) in pPGCs compared to soma and Epi.

(F) Immunofluorescence of GPR50 in E17 and E36 PGCs (indicated by yellow circles). PGCs are marked by SOX17 and Sda/GM2 in E17 and by DAZL in E36. Scale bar: 20µm.

**Figure S3**. Epigenetic reprogramming in pPGCs. Related to Figure 2 and Figure 3.

(A) Expression of DNMT3A and UHRF1 in E25 Gonadal pPGCs determined by IF. PGCs are marked by OCT4 (red). Scale bar: 20µm. Yellow circles indicate PGC.

(B) Immunofluorescence of 5hmC (top) and 5mC(bottom) in E36 Gonadal PGCs. PGCs are marked by Sda/GM2 (top, green) and SOX17 (bottom, red). Scale bar: 20µm. Yellow circles indicate PGC.

(C) Expression profile of selected components of PRC2 complex, Mll complex and chromatin remodellers in E11 Epiblast cells, E14 Somatic cells, E14 PGCs and E31 PGCs.

**Figure S4**. Extensive X chromosome Reactivation in Pre-migratory pPGCs. Related to Figure 4.

(A) Sum expression of (top) all Y-chromosome genes, (middle) single-copy Y-chromosome genes and (bottom) all X-chromosome genes.

(B) Bootstrap of X:A ratio of E14 somatic cells, E14 PGCs and E31 PGCs. Each dot represents one cell. P value: pairwise Wilcoxon test.

(C) Immunofluorescence of H3K27me3 in E14 PGC cluster. Yellow circle indicates PGC. Xi-associated H3K27me3 are detected in somatic cells (arrow) and some pPGCs (arrowhead). Scale bar: 20µm.

(D) Quantification of the number of H3K27me3 spots in E17 and E25 pPGC.

**Figure S5**. Level of methylation in Wk5 (E35) gonadal pPGCs revealed by BSSeq. Related to Figure 5.

(A) Violin plots showing CpG methylation levels at different repetitive elements.

(B) Expression profiles of major TE families in E11 epiblast, E14 somatic cells, E14 and E31 PGCs. p value. * p < 0.05; ** p < 0.01; *** p < 0.001; **** p < 0.0001 by pairwise Wilcoxon test.

(C) Distribution of major TE families that retain partial methylation (≥10%) in pig gonadal soma and PGCs. N: indicates the number of TEs that retain partial methylation, followed by the percentage of those partially methylated TEs among all TEs of the families indicated. L1 is overrepresented in pPGCs samples.

(D) Distribution of CpG methylation in non-TE and TE genomic tiles (800nt, with at least 5 CpG with 1x coverage).

See also Table S6.

**Figure S6**. DNA demethylation escapees show common and distinct features between mammals. Related to Figure 6.

(A) Enrichment analysis for GWAS catalogue terms showing disease and traits associated with conserved TE-poor escapee genes between pig and human.

(B) Enrichment analysis of GWAS catalogue terms for pig-specific TE-poor escapee genes.

(C) GO term enrichment analysis of genes overlapping demethylation-resistant pig-specific TE, Pre_0SS (top) and L1_SS (bottom).

(D) Enrichment analysis for SMART domains and tissue-specific expression (UniProt UP_TISSUE) of TE-poor escapee genes shared by at least two species (human, mouse, pig).

## Supplemental Tables

**Table S1**: List of samples used in this study.

**Table S2**: GO terms and KEGG enrichment analysis of DEGs.

**Table S3:** DEG comparison between pig and Cy monkey.

**Table S4**: Pluripotency signature gene sets.

**Table S5**: Sequencing statistics for PBAT.

**Table S6**: PGC-high TE repressors.

**Table S7**: Antibody List.

## STAR methods

### Resource availability

### Materials availability

This study did not generate new unique reagents

### Data and Code Availability

The scRNAseq and PBAT data generated under this study can be accessed from GEO: GSE155136.

### Experimental Model and Subject Details

#### Pig embryos and PGCs collection

All the procedures involving animals have been approved by the School of Biosciences Ethics Review Committee, The University of Nottingham. Embryos were retrieved from crossbred Large White and Landrace sows (2–3 years old) between days 11 to 35 after artificial insemination. E11 and E14 embryos were flushed from the uterine horns with warm washing buffer (PBS supplemented with 1% fetal bovine serum (FBS)). Later stage embryos (>E25) were manually dissected from the uterine horns and washed with washing buffer. Epiblast from E11 embryos were manually dissected and stored at -80°C before further processing for LC-MS (see below). PCR of AMEL gene was used for sex identification of E35 embryos before processed for FACS and PBAT library preparation (Sembon et al., 2008).

Pig PGC isolation was carried out as previously described (Hyldig et al., 2011a). Briefly, embryos between E14 to E35 were stored in DMEM/F-12 supplemented with 40% FBS at 4°C overnight before being processed the next day. Dissected posterior ends of E14 embryos containing PGC clusters and gonads from E31 and E35 embryos were digested at 37□°C for 30□mins using Collagenase IV (2mg/ml in DMEM), with gentle pipetting every 5 mins. The cell suspension was washed with DMEM, centrifuged and the pellet re-suspended in TrypLE Express (GIBCO) for further digestion at 37°C for 3-5 mins. Enzymatic digestion was neutralized with dissection medium (DMEM/F-12 with 10% FBS, 25 mM HEPES and 100 U/ml Penicillin-0.1 mg/ml Streptomycin). The cell suspension was filtered through a 40 µm cell strainer into FACS tube. Following centrifugation, cells were re-suspended and incubated in dissection medium with Sda/GM2 antibody (Klisch et al., 2011) for 30 mins at 4°C. After washing with dissection medium, cells were re-suspended and incubated in dissection medium with Alexa-488 Donkey Anti Mouse (Invitrogen) for 30 mins, and then diluted with dissection media and FACS sorted by MoFlo XDP Cell Sorter (Beckman Coulter). For PBAT, E35 Sda/GM2 + cells were sorted twice to ensure high purity.

#### Human embryonic tissues and collection of hPGCs

Human embryonic tissues were used under permission from NHS Research Ethical Committee, UK (REC Number: 96/085). Human embryonic samples were collected following medical or surgical termination of pregnancy carried out at Addenbrooke’s Hospital, Cambridge, UK with full consent from patients. Crown-rump length, anatomical features, including limb and digit development, was used to determine developmental stage of human embryos with reference to Carnegie staging (CS). The sex of embryos was determined by sex determination PCR as previously described (Bryja and Konecny, 2003).

Human embryonic genital ridges from two individual male embryos (developmental week 7-8, Carnegie stage 19) were dissected in PBS and separated from surrounding mesonephric tissues. The embryonic tissues were dissociated with 100 µl TrypLE Express (Life Technologies) at 37 °C for 30 minutes. Tissues were pipette up and down for ten times every 5 minutes to facilitate dissociation into single cell suspension. After that, samples were diluted with 100 µl FACS medium (PBS with 3% fetal calf serum & 5 mM EDTA) and centrifuged at 500 xg for 5 minutes. Cell pellet was suspended with FACS medium and incubated with 5 µl of Alexa Fluor 488-conjugated anti-alkaline phosphatase (AP) (BD Pharmingen, 561495) and 25 µl of PerCP-Cy5.5-conjugated anti-CD117 (BD Pharmingen 333950) antibodies for 15 minutes at room temperature with rotation at 10 revolutions per minutes (rpm) in dark. Cell suspension was then diluted in 1 ml FACS medium and centrifuged at 500 xg for 5 minutes. After removing the supernatant, the cell pellet was resuspended in FACS medium and passed through a 35µm cell strainer. Samples were subjected to FACS using the S3 Cell Sorter (Bio-Rad). hPGCs (AP- and CD117-positive) and the neighbouring gonadal somatic cells (AP- and CD117-negative) were collected and stored at -80 °C until mass spectrometry analysis.

#### Human ESC culture, hPGCLC induction and collection

Male hESCs with a NANOS3–tdTomato reporter was established previously (Kobayashi et al., 2017) and confirmed as mycoplasma negative. hESCs were maintained on vitronectin-coated plates in Essential 8 medium (Thermo Fisher Scientific) according to manufacturer’s protocol. Cells were passed every 3-5 days using 0.5 mM EDTA in PBS without breaking cell clumps.

hPGCLCs were generated using a two-step protocol as described before (Kobayashi et al., 2017). Briefly, trypsinized hESCs were seeded on vitronectin-coated dish at 200,000 cells per well in 12-well plate and cultured in mesendoderm induction medium for 12 hours. Mesendoderm medium consisted of aRB27 basal medium (Advanced RPMI 1640 Medium (Thermo Fisher Scientific) supplemented with 1% B27 supplement (Thermo Fisher Scientific), 0.1 mM NEAA, 100 U/ml penicillin, 0.1 mg/ml streptomycin, 2 mM L-glutamine), 100 ng/ml activin A (Department of Biochemistry, University of Cambridge), 3 μM GSK3i (Miltenyi Biotec) and 10 μM of ROCKi (Y-27632, Tocris Bioscience).

To induce hPGCLCs, pre-mesendoderm cells were trypsinized into single cells and harvested into Corning Costar Ultra-Low attachment multiwell 96-well plate (Sigma) at 4,000 cells per well in hPGCLC induction medium, which composed of aRB27 medium supplemented with 500 ng/ml BMP4,10 ng/ml human LIF (Department of Biochemistry), 100 ng/ml SCF (R&D systems), 50 ng/ml EGF (R&D Systems), 10 μM ROCKi, and 0.25% (v/v) poly-vinyl alcohol (Sigma). Cells were cultured as floating aggregate for 5 days. Aggregates were trypsinized with 0.25% trypsin/EDTA at 37 °C for 5-15 min. Cell suspension was subjected to FACS by SH800Z Cell Sorter (Sony). NANOS3–tdTomato-positive hPGCLCs and NANOS3–tdTomato-negative neighbouring cells were collected for mass spectrometry analysis.

### Method Details

#### Isolation of single cells for single-cell library preparation

FACS sorted cells were washed in a small drop of PBS-PVP and single cells were manually collected with thin capillaries and placed into PCR tubes to prepare single-cell cDNA libraries following the Smart-seq2 protocol (Picelli et al., 2014).

Briefly, single cells were lysed by incubation at 72□°C for 3□min in PCR tubes containing 4□μl of cell lysis buffer, oligo-dT primer and dNTP mix. Reverse transcription and PCR pre-amplification were carried out with SuperScript II (Invitrogen) and KAPA HiFi HotStart ReadyMix (KAPA Biosystems) respectively according to Picelli et al. protocol (Picelli et al., 2014). PCR products were purified using Ampure XP beads (Beckman Coulter), and library size distribution was checked on Agilent dsDNA High Sensitivity DNA chips on an Agilent 2100 Bioanalyzer (Agilent Technologies). Concentration was quantified using Qubit Quant-iT dsDNA High-Sensitivity Assay Kit (Invitrogen). Samples with more than 0.2□ng□µl^−1^, free of short fragments (<500□bp) and with a peak at around 1.5–2□kb were selected for library preparation with Nextera XT DNA Library Preparation Kit (Illumina). Tagmentation reaction and further PCR amplification for 12 cycles were carried out, and PCR products were again purified using Ampure XP beads. Quality of the final cDNA library was analysed on an Agilent high sensitivity DNA chip. Final cDNA libraries had an average size of 700–800□bp and were quantified using NEBNext Library Quant Kit for Illumina (New England BioLabs) following the manufacturer instructions. Finally, libraries were pooled in groups of 50 with a 2□nM final concentration, and DNA sequencing was performed on a HiSeq 2500 Sequencing System (Illumina).

#### Single-cell RNA-Seq data analysis

Raw PE reads were trimmed against adaptor sequences by scythe (v0.981), and quality-trimmed by sickle (v1.33) using default settings. Trimmed reads were directionally aligned to the pig genome (Sus scrofa v11) by hisat2 (v2.1.0) with *-know-splicestie-infile* setting to increase mapping accuracy of splicing reads. Uniquely and correctly mapped reads were extracted for the downstream analysis. htseq-count was used to count the number of reads aligned to each gene (Sus scrofa v11.2 ensembl annotation build 91). Gene expression level was calculated and normalised by Transcripts Per Kilobase Million (TPM).

Low quality cells were filtered out from the dataset to reduce the downstream analysis noise. First, the total number of reads mapped to gene transcripts was calculated for each cell, and those with less than 1 million were removed. Second, the proportion of reads aligned to mitochondrial genes was estimated, as a high proportion suggests poor quality cells (Ilicic et al., 2016). The proportion cut-off was set at 0.5. Only cells of proportions below 0.5 were kept for the next analysis. Third, 2 outlier cells were identified by t-SNE dimensionality reduction. A total of 14,873 out of 25,880 annotated genes were identified in at least 3 cells with TPM□>□1.

The R package “scater” was applied to normalise read counts of genes for each good quality cell with acceptable sequencing coverage. A non-linear approach, t-stochastic neighbour embedding (t-SNE), was used to identify the relations between cells using normalised read counts. Unsupervised hierarchical clustering using all expressed genes as input was conducted on all filtered cells by normalised read counts in log2 scale. The distance method was euclidean, and the cluster method was ward.D2.

#### Differential expression and enrichment analysis

Pairwise comparisons of single-cell differential expressions were performed by SCDE using normalised read counts among four embryo stages. Two-tailed adjusted p-value were calculated using cZ scores from Benjamini–Hochberg multiple testing corrections, which followed a normal distribution. Significantly expressed genes were selected with a p-value

<0.05 as the threshold. Euclidean distance and default hclust were applied to determine the relationships between cells and between genes. Gene Ontology (GO) gene set enrichment analysis with DEGs utilised goseq for each pairwise comparison, also with upregulated DEGs and downregulated DEGs separately. GO term annotation was retrieved from the Ensembl database (Sus scrofa v11.1 ensembl annotation version 91). Enrichment analysis of biological pathways (KEGG) was performed with DEGs by R package “clusterProfiler”. Ensembl gene IDs of DEGs were mapped to NCBI gene IDs for KEGG pathway prior to enrichment analysis.

#### Inference of embryonic sex

Expressions of all the single-copy genes on chrY were summed up to determine the gender of each cell. First, any cell with the total TPM of chrY single-copy genes ≥□10 was regarded a male cell. Others were regarded as female cells. Then, the ratios of the total gene expressions between chrY and chrX (∑ ChrY Total TPM / ∑ ChrX Total TPM) were calculated across all cells. Any pre-determined male cell with the ratio lower than the maximal ratio of pre-determined female cells was regarded as the female cell.

#### Chromosome X dosage compensation analysis

Genes of chromosome X and three autosomes (chr1, chr2, chr3) were extracted, and the geometric mean TPM of chromosomal expressed genes was calculated for each cell separately. Then the overall geometric mean TPM was obtained for each developmental stage by embryo sex, as well as the total TPM. Each TPM value was incremental by one (TPM□+□1) for the calculation of geometric mean TPM. Only shared expressed genes between female and male cells were taken into account in the calculation of female/male expression ratio for each chromosome. Median Female/Male expression ratio was estimated for each stage across the whole chromosome X with 1□Mb window. The ratio of chrX/auto in each cell was inferred by the median value of bootstrapped ratios. Each ratio was estimated by the total TPMs of a certain number of random-selected genes. The median ratios were grouped by embryo sex.

#### Analyses of allelic expression

Trimmed reads were aligned to chromosome X of the pig genome (Sus Scrofa v11.1) by hisat2. Duplicated reads were marked by picard (v2.12.1). GATK (v3.8) was used to retrieve allelic read counts for SNVs annotated in dbSNP. Only validated SNVs (dbSNP flag VLD) were extracted for downstream analysis. SnpEff (v4.3) was applied to annotate called SNVs with Sus scrofa v11.1 ensembl annotation. Low coverage SNVs (<3 reads) were excluded from the analysis, and we only kept SNVs that occurred at least in two different cells for each stage. The expressions of mono-/bi-allelic genes were inferred based on SNVs in each female cell of each stage.

#### Single cell trajectory analysis

Trajectory modelling and pseudotemporal ordering of cells was performed using TPM data with Monocle 2 (Qiu et al., 2017) (version 2.12.0). Top 1000 significant differentially expressed genes between clusters were used for ordering the cells.

#### Comparison of pig, human and cynomolgus monkey datasets

In total, dataset of E14-31 pig cells (128 from our study), processed data of Wk4-7 human cells (149) retrieved from GSE86146 (Li et al., 2017) and processed data of E13-55 cy monkey cells (100) retrieved from GEO: GSE76267, GSE74767 and GSE67259 (Sasaki et al., 2016) were included in the comparison. Natural log-transformed relative expression (i.e. ln(TPMs+1) in pig and human, log(RPMs+1) in cy monkey) of common genes (i.e. homologues genes with same gene name) across three species were imported and processed by *FindIntegrationAnchors* and *IntegrateData* (k.filters set as “NA”) functions in Seurat (version 3.1.2)(Stuart et al., 2019). Dimensionality reduction by UMAP was then performed for the integrated dataset.

Expression of selected lineage markers and epigenetic modifiers in E14-31 pig cells, Wk4-7 human cells and E13-55 *Cynomolgus* cells were plotted separately with pheatmap package.

#### Cell cycle analysis

Default settings of *CellCycleScoring* function in Seurat were used to score the cell cycle phases of each single cell. In brief, single cells were assigned a score with *AddModuleScore* function based on its expression of G2/M- and S-phase markers provided in Seurat. The single cells highly expressing G2/M- or S-phase markers were assigned as G2/M- or S-phase cells, respectively, and the single cells not expressing any of the two categories of genes were assigned as G1 phase.

#### Signature set analysis

With the processed single cell RNA-seq data of pig embryos from Ramos-Ibeas et al. (Ramos-Ibeas et al., 2019) we used *FindMarkers* function in Seurat (Wilcoxon rank sum test) to identify the highly expressed genes (avg_logFC >=1 and adjusted.p <=0.05) as the signature set in E6 ICM, E8 epiblast and E11 epiblast. Next, we calculated the relative average expression level of each signature set with *AddModuleScore* function of Seurat in single cells of E14 Soma, E14 PGC and E31 PGC, which was then visualized by heatmap using pheatmap package.

#### PBAT library construction

PBAT libraries were prepared as described previously (Tang et al., 2015) with some modifications. The Sda/GM2-positive (PGCs) and -negative (Somatic) cells collected by FACS were lysed with lysis buffer (0.1% SDS, 50 ng/ml carrier RNA (QIAGEN) and 1 mg/ml proteinase K (Zymo Research) in DNase-free water) for 60 min at 37°C. Unmethylated lambda phage DNA (0.2 ng/sample) (Promega) was spiked into the sample before bisulfite treatment with the Methylcode Bisulfite Conversion Kit (Invitrogen) according to the manufacturer’s instructions, except that the bisulfite conversion step was increased to 3.5 hours. Bisulfite-treated DNA was re-annealed to double-stranded DNA using Klenow fragments (3’–5’ exo-) (New England Biolabs) with a 5’ biotin tagged primer consisted of an Illumina adaptor followed by 6 random nucleotides at the 3’ end (BioPEA2N4: 5’-[btn] CTACACGACGCTCTTCCGATCTNNNNNN-3’)(Clark et al., 2017).

The biotinylated first strand molecules were captured using Dynabeads M280 Streptavidin (Invitrogen) and then reannealed to double-stranded DNA again using Klenow fragments (3’–5’ exo-) with random primers containing Illumina adaptors (Rev_N6_PE: 5’-TGCTGAACCGCTCTTCCGATCTNNNNNN-3’)(Clark et al., 2017).

Template DNA strands were then synthesized as cDNA with a second strand (where unmethylated C’s were converted to T’s) and then amplified with 11 cycles using KAPA HiFi HotStart Readymix (Roche) with the Illumina primer PE 1.0 (5’-AATGATACGGCGACCACCGAGATCTACACTCTTTCCCTACACGACGCTCTTCCGATC*T-3’) and iPCRTag (5’-CAAGCAGAAGACGGCATACGAGATAACGTGATGAGATCGGTCTCGGCATTCCTGCTGA ACCGCTCTTCCGATC*T-3’).

Size fractionation was performed on the eluted DNA with Agencourt AMPure XP (Beckman Coulter). Concentrations of PBAT libraries were determined by qPCR using NEBNext Library Quant kit (NEB). Libraries were subjected to paired-read 150bp sequencing on HiSeq 4000 sequencing system (Illumina). Coverage information was summarized in Table S5.

#### DNA methylation analysis

The quality of raw reads was determined by FastQC to ensure that the experimental setup and sequencing were successful. Raw reads were trimmed by skewer first to remove adaptor sequences and reads with low sequencing qualities (Jiang et al., 2014). Then, both the ends of paired-end reads were trimmed to improve the mapping efficiency. Forward reads were trimmed by 10 bases at the beginning, while reverse reads were trimmed by 5 bases at the end.

Trimmed reads were directionally aligned against the pig genome (Sus Scrofa v11.1) in the paired-end mode by hisat2 using Bismark pipeline with *--pbat*. --score_min was L,0,-0.4. *deduplicate_bismark* was applied to remove the potential PCR duplicates with default settings (Krueger and Andrews, 2011). Unmapped reads were re-aligned with the same parameters in the single-end non-direction mode to rescue misaligned paired-end reads due to the incorrect insert size resulting from the narrow sequencing area. The single-end alignment was merged with the paired-end alignment after deduplication.

To compare the pig PBAT datasets with those from human (Tang et al., 2015) and mouse (Kobayashi et al., 2013), reads were trimmed up to 100 nt for all three species, and were mapped via single-end only and sampled to the same depth.

The detection of methylated cytosines was done by *bismark_methylation_extractor*, which can provide the genome-wide cytosine methylation status. The spike-in unmethylated lambda phage DNA was also included in the analysis to examine the efficiency of bisulphite conversion in the samples.

The annotation of the methylation level was calculated by the module *roimethstat* of *MethPipe* according to the locations of CpG islands and CGI shores, the genomic features and by the repeat density (Song et al., 2013). Annotations of CpG islands, genes, promoters and repeat regions were downloaded from UCSC and Ensembl databases. Promoter regions were defined as sequences located between 1,000 bp upstream and 500 bp downstream of a transcription start site. Promoters with high-CpG content (HCP) contain a 500 bp region with a CpG ratio larger than 0.75 and a GC content larger than 55 %. Promoters with low-CpG content (LCP) do not contain a 500-bp region with a CpG ratio larger than 0.48. Intermediate-CpG promoters (ICPs) are neither HCP nor LCP.

Hypermethylated regions (HyperMR) were identified by the *hmr* function of *MethPipe*. Escapees were defined as regions which have more than 20% of CpGs with >= 5x with at least 30% methylation level in human and 15% in pig and mouse. TE-poor escapees were defined as less than 10% of regions overlapped with repeats. TE-rich escapees were defined as more than 10% of regions overlapped with repeats.

#### TE Expression Analysis

Repeat regions were downloaded from UCSC database including all the sub families. *featureCounts* (Liao et al., 2014) was used to determine the number of reads aligned to each region with *-M* option. Expression level was calculated and normalised by Reads Per Kilobase Million (RPM).

#### Immunofluorescence staining of porcine tissues

Embryos were processed as previously described (Kobayashi et al., 2017). Briefly, embryos and gonads were fixed in 4% paraformaldehyde (PFA)/PBS overnight (ON) at 4°C. Fixed embryos were incubated in 30% sucrose/PBS for two days at 4°C prior to mounting in optimal cutting temperature (OCT) compound. Cryosections were cut at 5-7 µm onto Superfrost plus glass slides. Sections were left to air dry for 1-2 h before IF.

For IF, cryosections were washed with PBS for 10 mins to remove OCT compound. Antigen retrieval was then performed by boiling the slides in 0.01M Citrate Buffer (pH 6.0) for 10 min. Sections were permeabilized with 1% Triton-X100 in PBS for 15 min. Triton-X100 was washed three times for 5 min each, and blocking solution (PBS supplemented with 5% BSA and 10% Donkey serum) was added for 1.5 h. After blocking, sections were incubated with the desired primary antibody (Table S7) ON at 4°C in a humidified chamber. Slides were then washed three times with 0.1% Tween-20/PBS. Slides were then incubated with fluorescent (Alexa Fluorophore 488, 555, and/or 647; Invitrogen)-conjugated secondary antibodies for 40 min at room temperature (RT). Slides were mounted with Fluoroshield with DAPI (Sigma) and sealed with nail varnish. Slides were kept at -20 °C until observed.

Image acquisition was performed using SimplePCI capture software on an epifluorescence microscope (Leica). Fiji was used for cell count and fluorescence quantification of ROI (Schindelin et al., 2012). For fluorescence quantification, background intensity was subtracted to generate corrected total cell fluorescence (CTCF), i.e. CTCF = Integrated Density – (Area of selected cell X Mean fluorescence of background readings)(McCloy et al., 2014).

#### Mass spectrometry

Genomic DNA from E11 epiblast and FACS-sorted pPGCs was extracted using Quick-DNA/RNA Miniprep kit (Zymo Reasearch) following the manufacturer’s instructions and eluted in LC–MS grade water. DNA was digested to nucleosides using a using a nucleoside digestion mix (NEB). The nucleosides were separated on an RRHD Eclipse Plus C18 2.1 × 100 mm 1.8u column using the HPLC 1290 system (Agilent) and mobile phases 100% water 0.1% formic acids and 80% methanol, 0.1% formic acids. Quantification was carried out in an Agilent 6490 triple quadrupole mass spectrometer on multiple reaction monitoring mode (MRM). To calculate the concentrations of individual nucleosides, standard curves were generated (dC and dG from Berry and Associated; 5mdC and 5hmdC from CarboSynth). All samples and standard curve points were spiked with a similar amount of isotope-labelled synthetic nucleosides (13C15N-dC and 13C15N-dG purchased from Silantes, and d3-mdC and d215N2-mhdC was obtained from T. Carell (Center for Integrated Protein Science at the Department of Chemistry, Ludwig-Maximilians-Universität München, Germany). The threshold for quantification is a signal-to-noise above ten (calculated with a peak-to-peak method). Limit of quantification (LOQ) was 0.025 fmol for 5mdC and 5hmdC, and 0.5 fmol for dC and dG.

#### Statistical analysis

Statistical differences in 5hmC and 5mC levels determined by LC–MS, were determined with ANOVA and Holm’s post hoc test. Differences in female to male expression ratio across X chromosome and X:A ratio in E14 PGC, E31 PGC and E14 Somatic cells, were calculated using pairwise Wilcoxon test. To evaluate the statistical differences in number of biallelically expressed genes in E14 PGC, E31 PGC and E14 Somatic cells, p value is determined by Kruskal-Wallis test followed by Dunn’s test. Statistical differences in *KDM6A* expression in E14 cells, *was calculated* using Mann-Whitney U-test. Differences in H3K27me3 quantification in E17 migratory PGC and surrounding somatic cells, were calculated with Mann-Whitney U-test. Differences in expression profiles of major TE families in E11 epiblast, E14 somatic cells, E14 and E31 PGCs were calculated using pairwise Wilcoxon test.

## Notes

### Competing Interest Statement

The authors have declared no competing interest.

