## Supplementary material for "Specification and epigenetic resetting of the pig germline exhibit conservation with the human lineage": Suppl. figures

### Figure S1

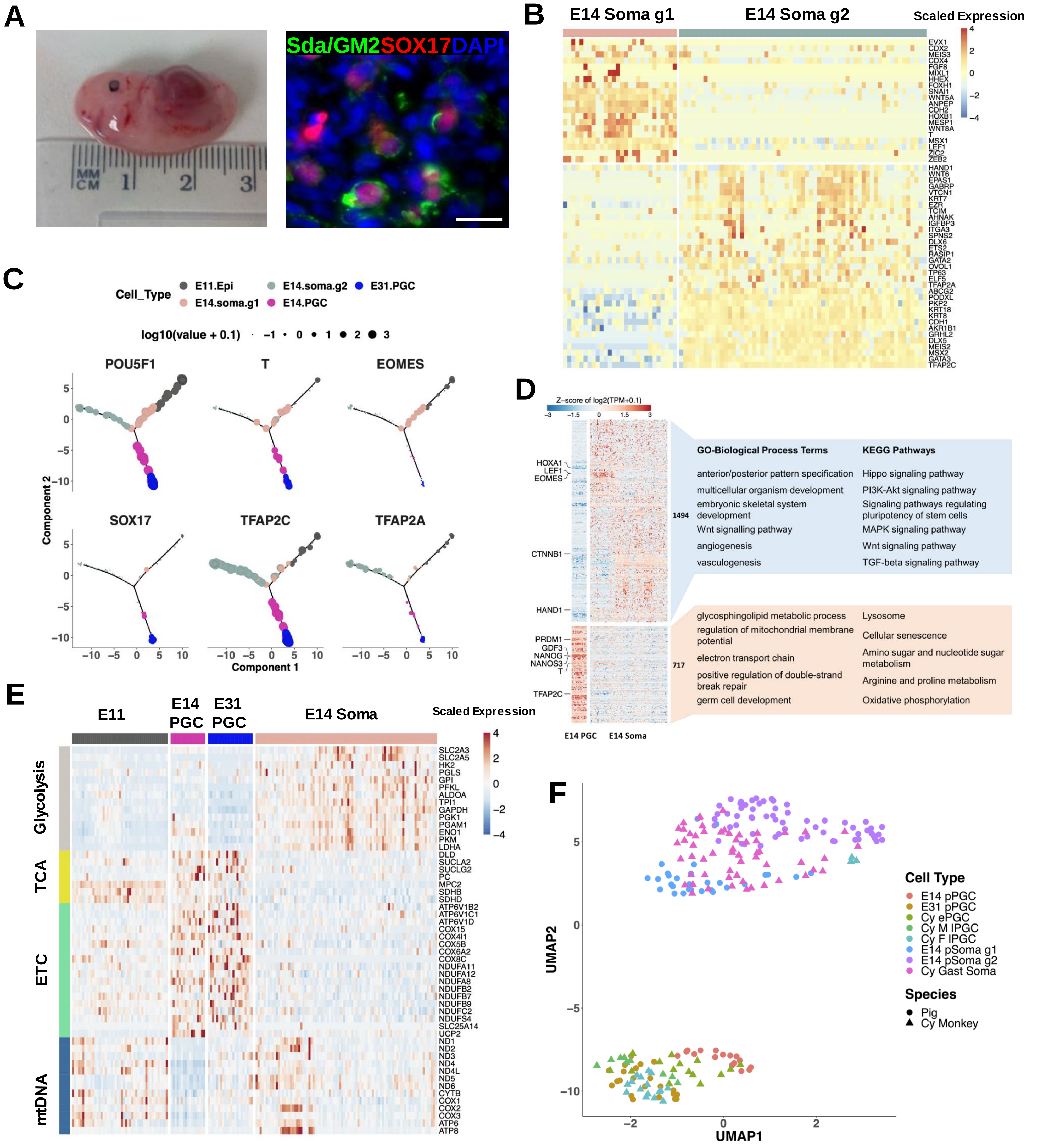


**Figure S1.** Gene expression differences between pPGC and surrounding cells. Related to Figure 1.

(A) Isolation of PGCs from from E31 embryos using the cell surface marker Sda/GM2 with FACS. A section of a fetal ovary shows PGCs stained for the indicated markers (right). Scale bar: 20µm.
(B) Expression heatmap of DEGs in somatic cells found in the posterior end of E14 embryos.
(C) Expression of selected lineage markers on trajectory of single cells.
(D) Gene expression heatmap, GO terms and KEGG pathways for DEGs between E14 PGCs and somatic cells. See also Table S2.
(E) Gene expression heatmap of cellular metabolism and mitochondrial DEGs in different cells types. ETC: electron transport chain. mtDNA: mtDNA-encoded components. TCA: tricarboxylic cycle.
(F) UMAP plot showing integration of *Cynomolgus* (E13-55) and pig (E14-31) PGCs and respective somatic cells.

### Figure S2

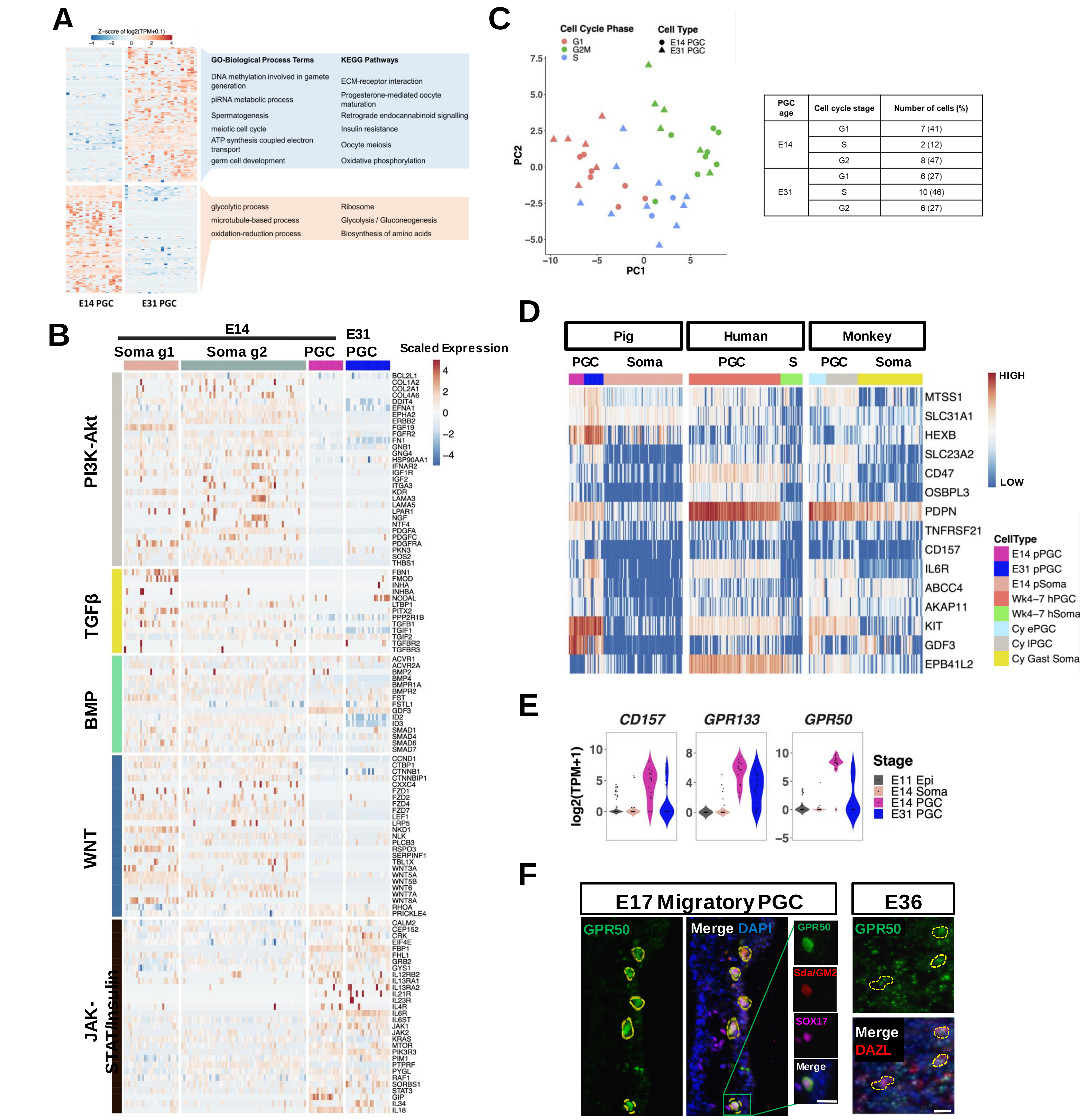


| PGC age | Cell cycle stage | Number of cells (%) |
|---|---|---|
| E14 | G1 | 7 (41) |
| | S | 2 (12) |
| | G2 | 8 (47) |
| E31 | G1 | 6 (27) |
| | S | 10 (46) |
| | G2 | 6 (27) |

**Figure S2**. Molecular differences between E14 and E31 pPGC, Related to Figure 1.

(A) Gene expression heatmap, GO terms and KEGG pathways for DEGs between E14 and E31 PGCs. See also Table S2.
(B) Heatmap of selected genes from different signalling pathways expressed in E14 and E31 cells.
(C) Determination of cell cycle stage based on relative average expression (module score) of canonical cell cycle markers in E14 and E31 PGCs.
(D) Expression heatmap of cell surface and membrane proteins in hPGCs, cyPGCs and pPGCs compared to somatic cells.
(E) Violin plots showing expression of selected cell surface proteins in pPGCs compared to soma and Epi.
(F) Immunofluorescence of GPR50 in E17 and E36 PGCs (indicated by yellow circles). PGCs are marked by SOX17 and Sda/GM2 in E17 and by DAZL in E36. Scale bar: 20µm. .

### Figure S3

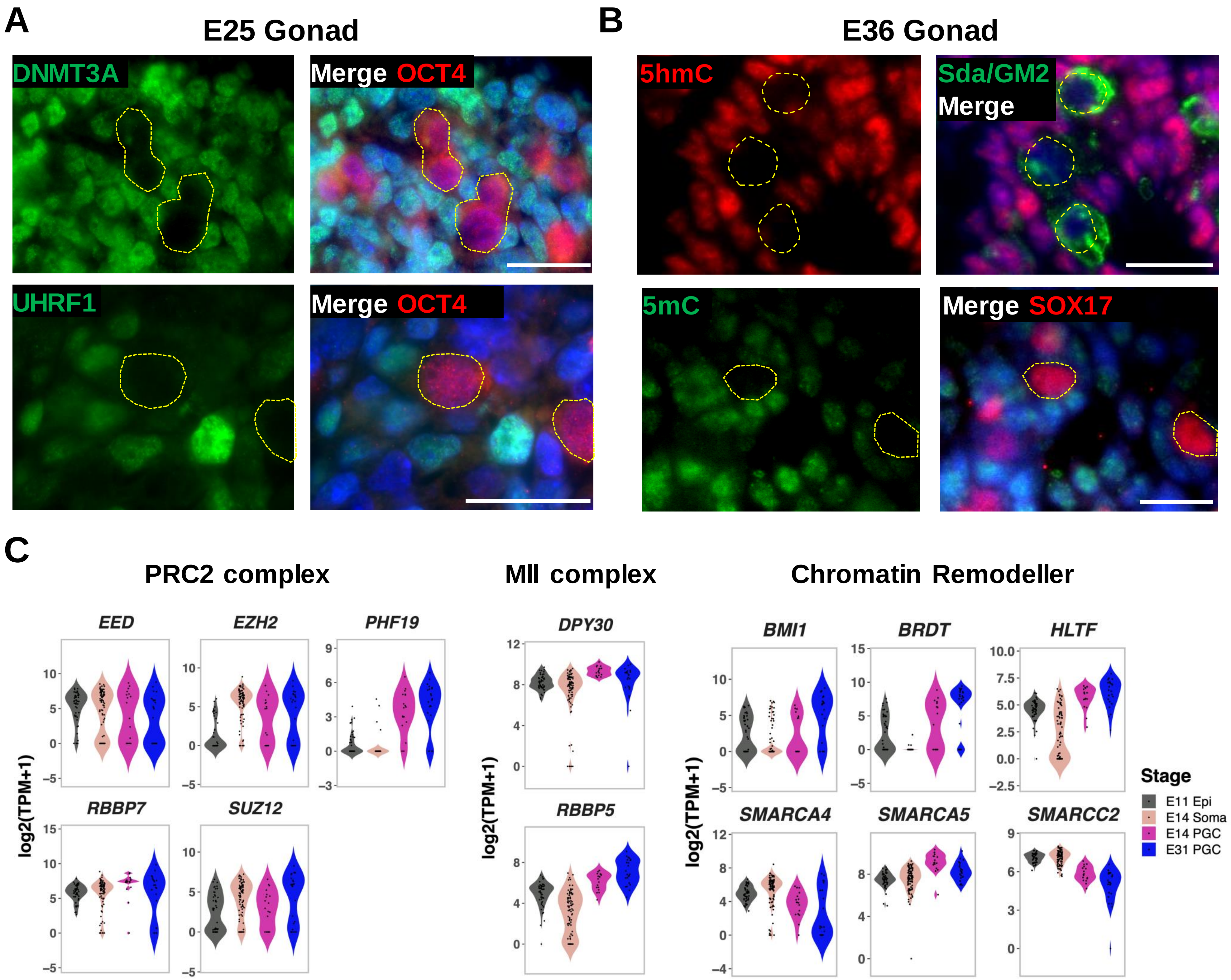


**Figure S3**. Epigenetic reprogramming in pPGCs. Related to Figure 2 and Figure 3.

(A) Expression of DNMT3A and UHRF1 in E25 Gonadal pPGCs determined by IF. PGCs are marked by OCT4 (red). Scale bar: 20µm. Yellow circles indicate PGC.
(B) Immunofluorescence of 5hmC (top) and 5mC(bottom) in E36 Gonadal PGCs. PGCs are marked by Sda/GM2 (top, green) and SOX17 (bottom, red). Scale bar: 20µm. Yellow circles indicate PGC.
(C) Expression profile of selected components of PRC2 complex, Mll complex and chromatin remodellers in E11 Epiblast cells, E14 Somatic cells, E14 PGCs and E31 PGCs.

### Figure S4

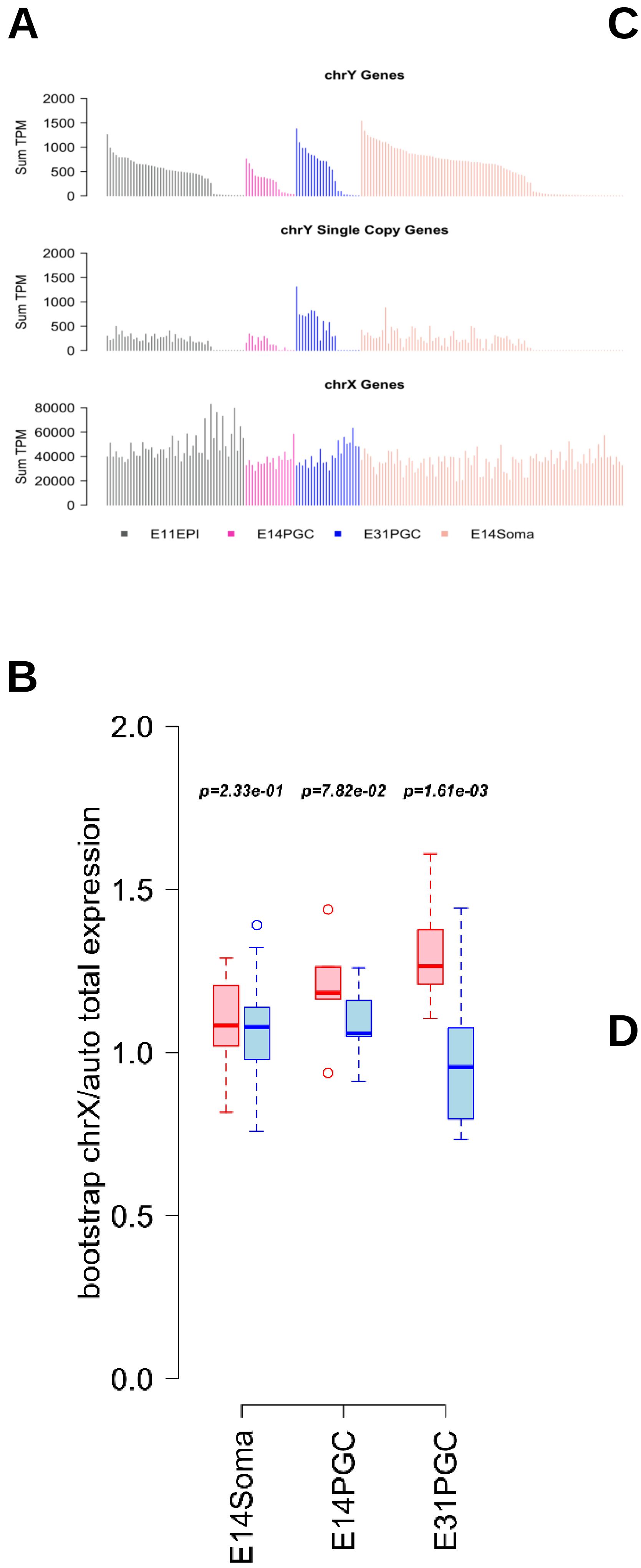


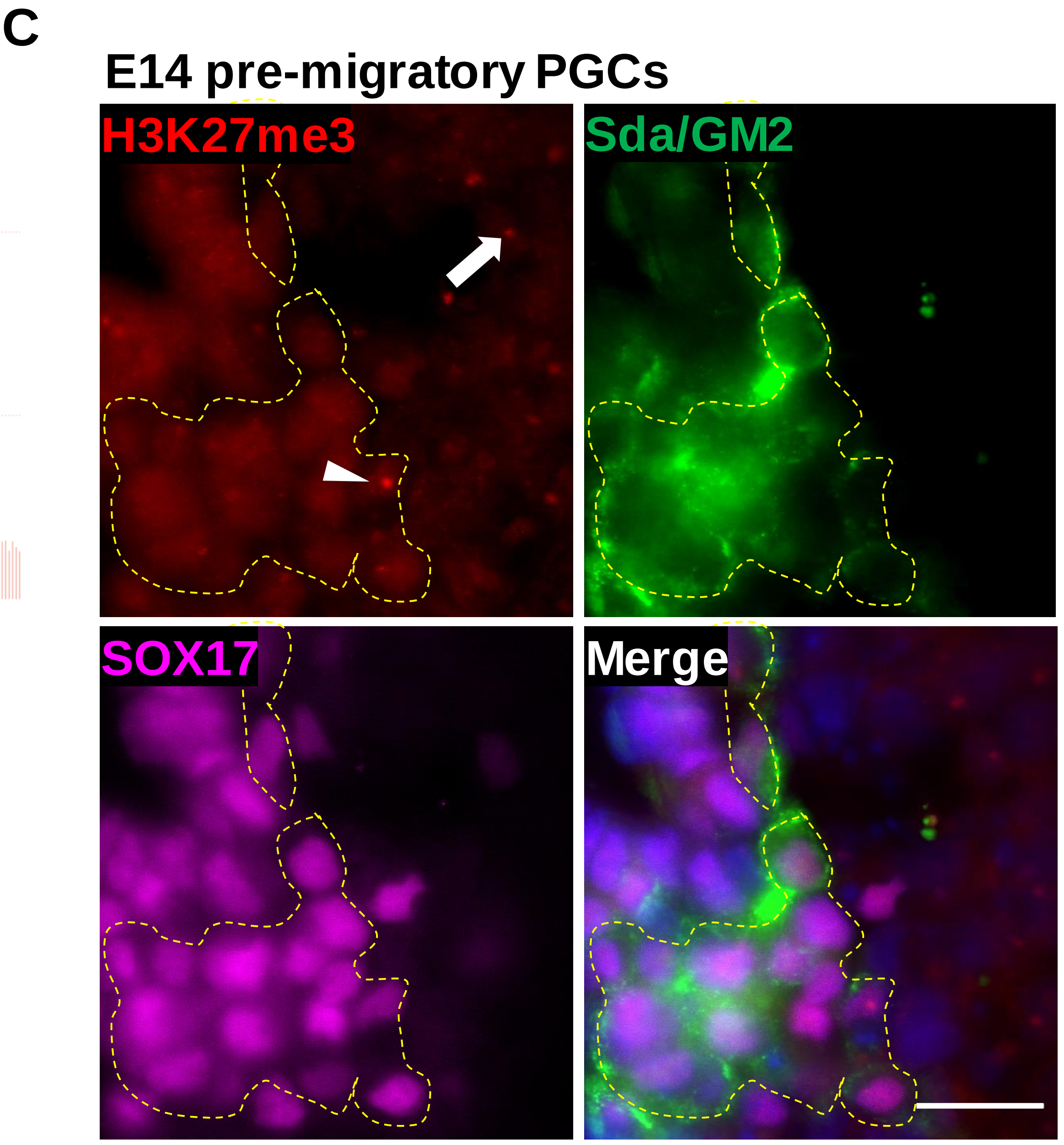


**D**

| | No H3K27me3 spot | Strong H3K27me3 spot | Ambiguous |
|---|---|---|---|
| **No. of E17 PGC (%)** | 96 (72) | 19 (14.5) | 18 (13.5) |
| **No. of E25 PGC (%)** | 122 (73) | 12 (7.2) | 33 (19.8) |

**Figure S4**. Extensive X chromosome Reactivation in Pre-migratory pPGCs. Related to Figure 4.

(A) Sum expression of (top) all Y-chromosome genes, (middle) single-copy Y-chromosome genes and (bottom) all X-chromosome genes.
(B) Bootstrap of X:A ratio of E14 somatic cells, E14 PGCs and E31 PGCs. Each dot represents one cell. P value: pairwise Wilcoxon test.
(C) Immunofluorescence of H3K27me3 in E14 PGC cluster. Yellow circle indicates PGC. Xi-associated H3K27me3 are detected in somatic cells (arrow) and some pPGCs (arrowhead). Scale bar: 20µm.
(D) Quantification of the number of H3K27me3 spots in E17 and E25 pPGC.

### Figure S5

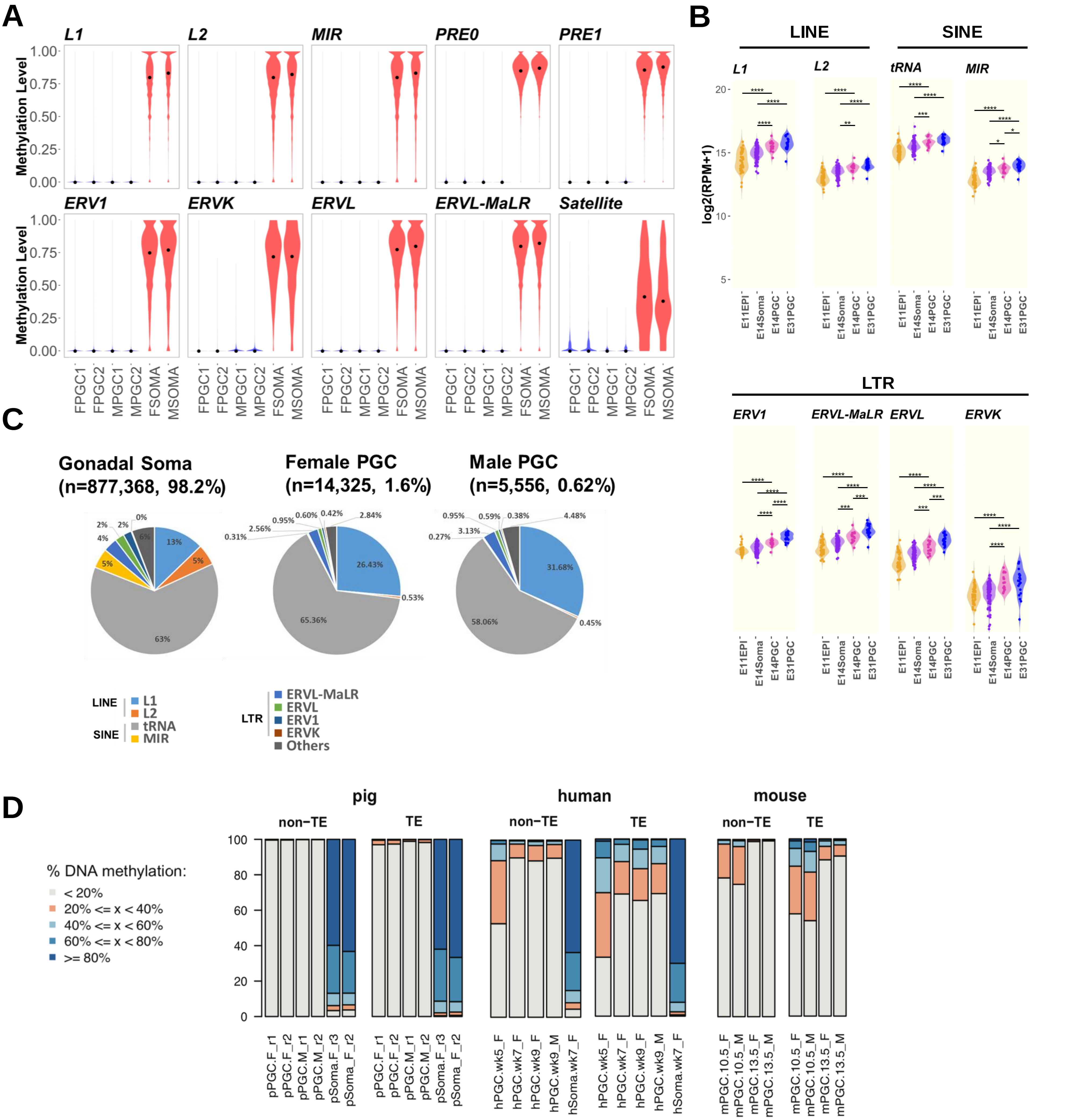


**Figure S5**. Level of methylation in Wk5 (E35) gonadal pPGCs revealed by BSSeq. Related to Figure 5.

(A) Violin plots showing CpG methylation levels at different repetitive elements.
(B) Expression profiles of major TE families in E11 epiblast, E14 somatic cells, E14 and E31 PGCs. p value. * $p < 0.05$; ** $p < 0.01$; *** $p < 0.001$; **** $p < 0.0001$ by pairwise Wilcoxon test.
(C) Distribution of major TE families that retain partial methylation (≥10%) in pig gonadal soma and PGCs. N: indicates the number of TEs that retain partial methylation, followed by the percentage of those partially methylated TEs among all TEs of the families indicated. L1 is overrepresented in pPGCs samples.
(D) Distribution of CpG methylation in non-TE and TE genomic tiles (800nt, with at least 5 CpG with 1x coverage).
See also Table S5.

### Figure S6

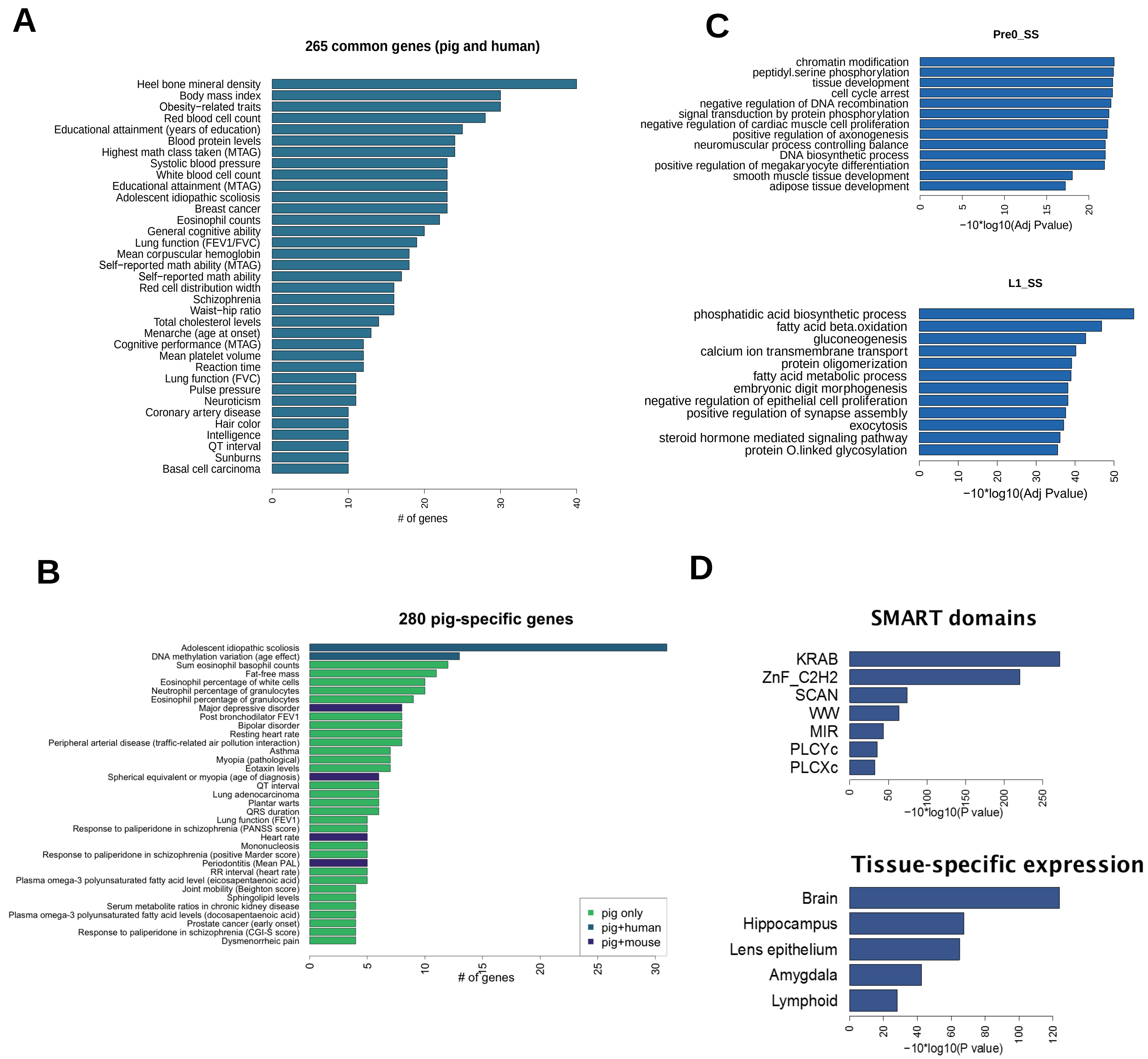


**Figure S6**. DNA demethylation escapees show common and distinct features between mammals. Related to Figure 6.

(A) Enrichment analysis for GWAS catalogue terms showing disease and traits associated with conserved TE-poor escapee genes between pig and human.
(B) Enrichment analysis of GWAS catalogue terms for pig-specific TE-poor escapee genes.
(C) GO term enrichment analysis of genes overlapping demethylation-resistant pig-specific TE, Pre_0SS (top) and L1_SS (bottom).
(D) Enrichment analysis for SMART domains and tissue-specific expression (UniProt UP_TISSUE) of TE-poor escapee genes shared by at least two species (human, mouse, pig).
